# Dual control of lysogeny and phage defense by a phosphorylation-based toxin/antitoxin system

**DOI:** 10.1101/2022.09.05.506569

**Authors:** Yunxue Guo, Kaihao Tang, Brandon Sit, Jiayu Gu, Ran Chen, Jianzhong Lin, Shituan Lin, Xiaoxiao Liu, Weiquan Wang, Xinyu Gao, Zhaolong Nie, Tianlang Liu, Matthew K. Waldor, Xiaoxue Wang

## Abstract

Regulatory systems that maintain prophage quiescence integrate phage and host gene expression with environmental conditions^1,2^. In the opportunistic bacterial pathogen *Pseudomonas aeruginosa*, Pf filamentous bacteriophages play critical roles in biofilm formation and virulence^3-5^, but mechanisms governing Pf prophage activation in biofilms are largely unknown. Here, we report a new type of prophage regulatory module in a widely-distributed *P. aeruginosa* lineage that not only controls virion production of co-resident Pf prophages, but also mediates defense against diverse lytic phages. By comparing two lineages of the prototype *P. aeruginosa* strain PAO1 that harbor different Pf prophages, we identified a prophage-encoded kinase-kinase-phosphatase (KKP) system that controls Pf production in biofilms. KKP components exhibit dynamic stoichiometry, where high kinase levels in planktonic conditions maintain phosphorylation of the host H-NS protein MvaU, repressing prophage activation. During biofilm formation, phosphatase expression is heightened, leading to MvaU dephosphorylation and alleviating repression of prophage gene expression. KKP clusters are present in hundreds of diverse temperate prophages and other mobile elements across Gram-negative bacteria. Characterization of KKP modules from different species revealed that, in addition to regulating Pf phage lysogeny, KKP functions as a tripartite toxin-antitoxin system that mediates host defense from predatory lytic phages. KKP represents a new phosphorylation-based mechanism for prophage regulation and for phage defense. The dual function of this module raises the question of whether other newly described phage defense systems^6-9^ also regulate intrinsic prophage biology in diverse hosts.

---

Filamentous bacteriophages (inoviruses) infect diverse bacterial hosts^10,11^, including the opportunistic human pathogen *Pseudomonas aeruginosa*^12^. The filamentous ‘Pf’ phages in *P. aeruginosa* play important roles in virulence^5^ and formation of characteristic *P. aeruginosa* biofilms^3,4^, where expression of genes enabling virion production is markedly upregulated^13^. Pf virions directly participate in the formation of viscous liquid crystals^14-16^ and can suppress mammalian immunity by directly inhibiting phagocytosis and bacterial clearance^17^. Several host factors, such as H-NS-like nucleoid binding proteins, regulate the lysogeny of the Pf4 prophage^18-20^, but the mechanisms that control Pf prophage activation are largely uncharted. Pfs often encode conserved structural genes as well as disparate accessory genes whose functions are not clear^21-25^, suggesting genetic diversity within the Pf clade may have consequences for understanding Pf prophage biology and regulation. Additionally, although many *P. aeruginosa* strains encode multiple Pf prophages, interactions between co-resident Pf prophages have not been documented.

## Divergent Pf production in two globally-distributed PAO1 lineages

During studies of *P. aeruginosa* biofilm formation in a flow cell-based catheter system (Fig. 1a) that mimics chronic infection^26^, we observed that two major lineages of the prototypical *P. aeruginosa* strain PAO1 produced dramatically different titers of Pf virions in biofilms (Fig. 1b). These lineages were designated as ‘PAO1’, the original strain isolated in 1955^27^, and ‘MPAO1’, a widely distributed derivative of PAO1^28^. In the flow cell system, PAO1 and MPAO1 both formed visible thick, crystal violet-positive biofilms on the inner surface of the catheter after 3 days of growth (Fig. 1a). However, biofilm effluents from the two strains contained marked differences in phage titers. TEM of MPAO1 effluents at day 6 revealed characteristic bundles of phages with a wavy filamentous structure (length 2∼3 μm) (Fig. 1c), whereas no virions were detected in the PAO1 biofilm effluent. To visualize in situ Pf production in biofilms, the major coat protein (pVIII) of the Pf4 prophage, which is present in both MPAO1 and PAO1, was fused in-frame with GFP, creating a reporter of Pf4 production (Fig. 1d). Consistent with the PFU and TEM data, Pf4 production was readily observed in ridge-like structures within MPAO1 biofilms, but was only rarely detected in PAO1 biofilms (Fig. 1d).

**Fig. 1.**
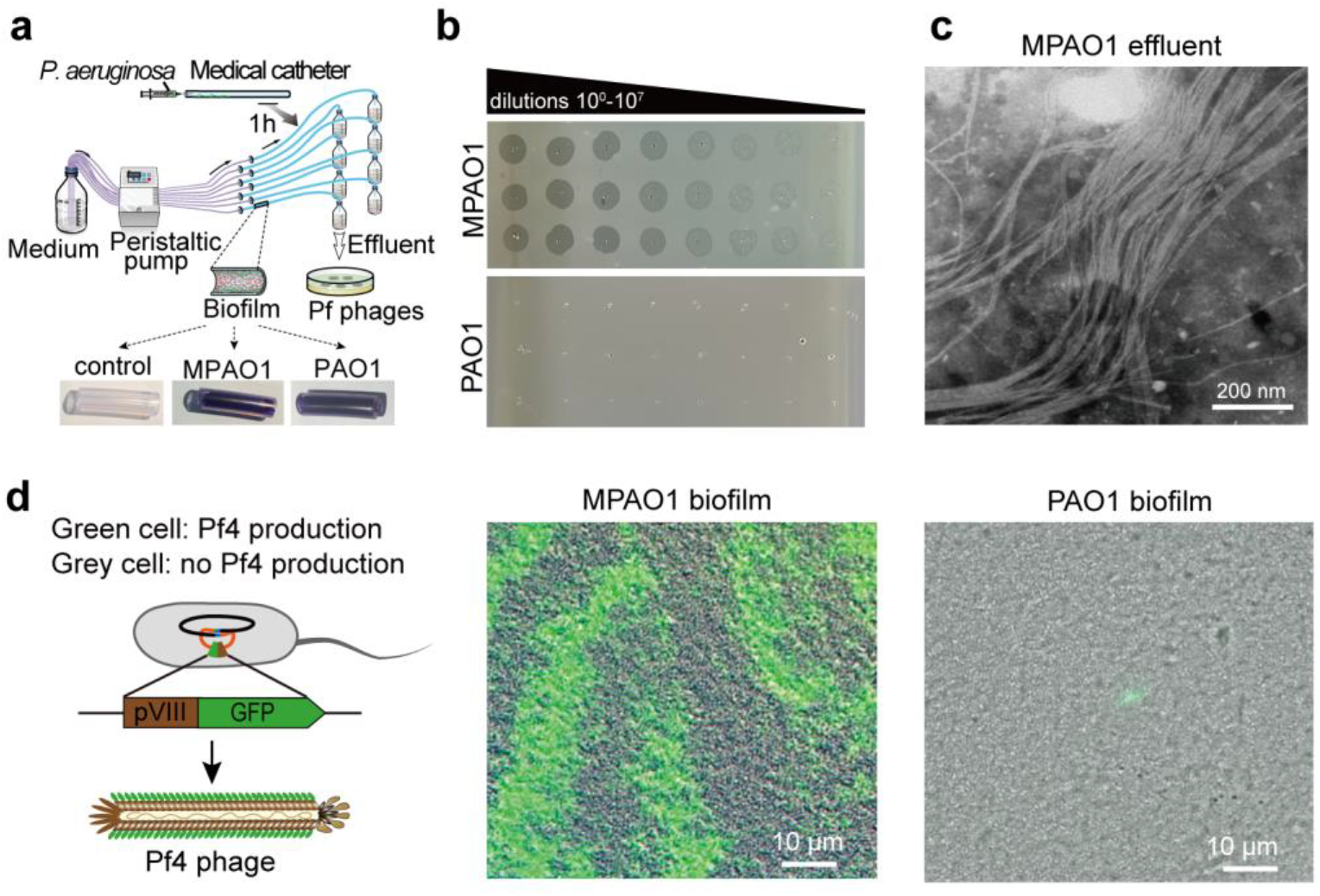
MPAO1, but not PAO1 produces Pf phages in biofilms. **a**, Schematic representation of the flow cell biofilm system. Representative biofilms stained with crystal violet are shown. **b**, Phage titers in day 6 biofilm effluents. **c**, Transmission electron microscopy (TEM) of biofilm effluents from day 6 MPAO1 biofilms. **d**, Fluorescence microscopy of day 6 biofilms in MPAO1 (left) or PAO1 (right) strains expressing the GFP tagged Pf4. At least three independent cultures were used for **b**-**d** and representative images are shown in **b**-**d**.

Genomic comparisons of PAO1 and MPAO1 have revealed minor genetic differences between these *P. aeruginosa* sublines that likely impact important phenotypes such as pyocyanin and pyoverdine production and virulence in animal models^29,30^. The largest difference between MPAO1 and PAO1 is the presence of a second Pf prophage, Pf6, in MPAO1. Whole-genome sequencing confirmed that PAO1 only harbors the Pf4 prophage, whereas MPAO1 harbors both the Pf4 and Pf6 prophage (Fig. 2a; Table S1). The Pf4 prophages in MPAO1 and in PAO1 are nearly identical, differing only in an 18 bp deletion near the *attL* site in MPAO1. The Pf6 prophage (12,150 bp) in MPAO1 is inserted in the tRNA^Met^ gene and the core region of the prophage, which encodes the virion capsid, assembly, and replication-related proteins (PA0717-PA0727), is highly similar (>98% nucleotide identity) to the core region of Pf4.

**Fig. 2.**
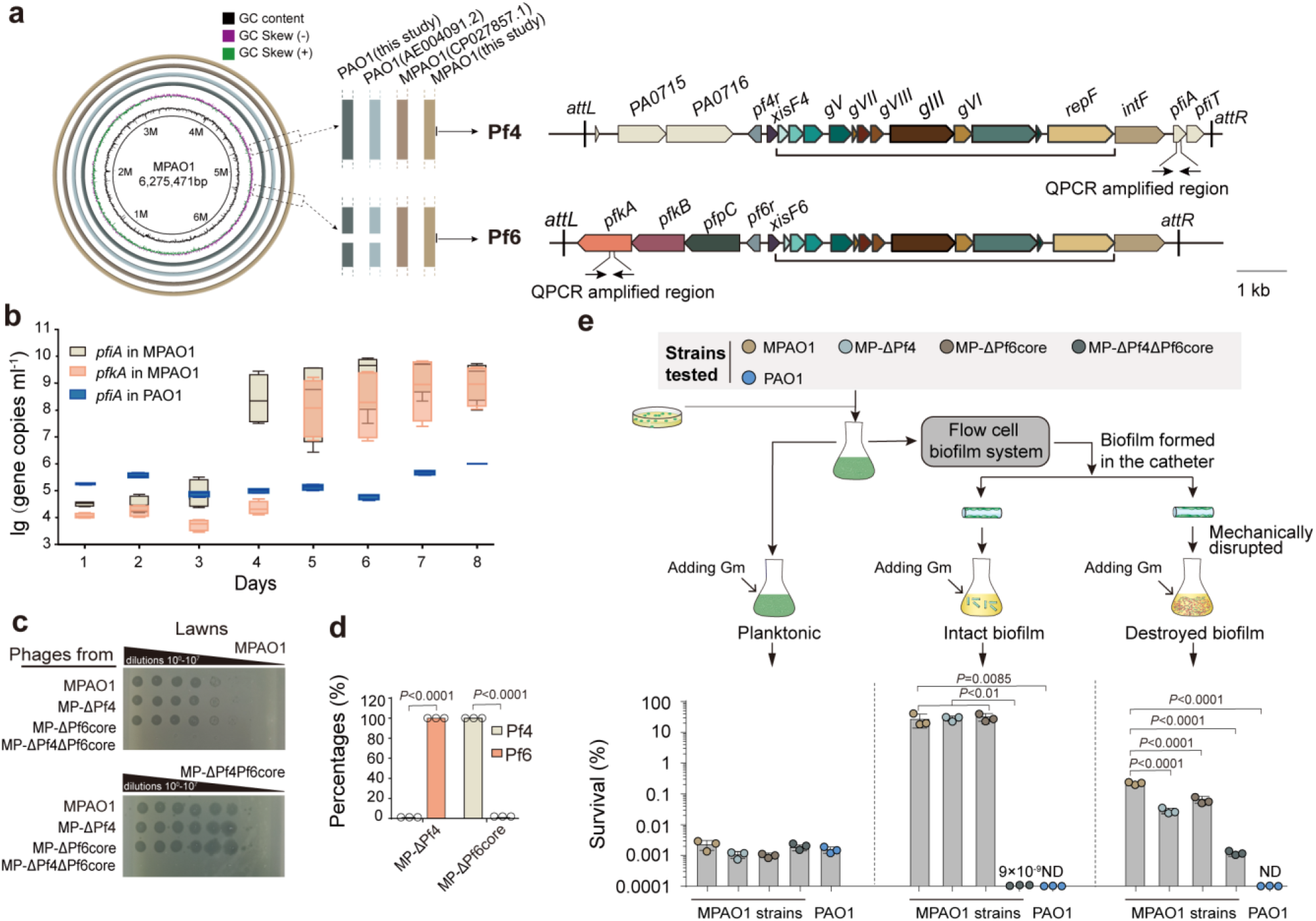
Pf4 and Pf6 are coordinately produced in MPAO1 biofilms and contribute to antibiotic resistance. **a**, Comparative genomic analysis of *de novo* sequenced MPAO1 and PAO1 with the first PAO1 and MPAO1 genomes deposited in NCBI. The MPAO1 sequenced here was used as reference for the annotation of Pf4 and Pf6 prophages with core Pf regions underlined. **b**, qPCR of Pf4-(*pfiA*) and Pf6-(*pfkA*) specific genes (shown in **Fig. 2a)** in MPAO1 and PAO1 biofilm effluents. **c**, Phage titers of day 6 biofilm effluents from the indicated MPAO1 strains plated on lawns of the two different host strains. **d**, Percentages of Pf4 and Pf6 phages in biofilm effluents from the MPAO1 mutant strains quantified by qPCR. **e**, Gentamicin (Gm) sensitivity of various strains in varying biofilm states. Experimental design is shown in top panel. ND indicates not determined. At least three independent cultures were used for **b-e** and representative images are shown in **c**.

Since MPAO1 harbors Pf4 and Pf6, we asked which prophage contributes to Pf virion production in *P. aeruginosa* biofilms. To independently monitor Pf4 and Pf6 phage DNA in biofilm effluents, the amounts of Pf4-(*pfiA*) and Pf6-(*pfkA*) specific genes were measured by qPCR (Fig. 2a). The abundance of both *pfiA* and *pfkA* increased over time (Fig. 2b), suggesting that production of both Pf4 and Pf6 is activated in MPAO1 biofilms. In contrast, levels of *pfiA* were unchanged in PAO1 biofilms, confirming that Pf4 is not produced in this lineage (Fig. 2b). An MPAO1 strain deleted for the Pf4 prophage (MP-ΔPf4) was generated to test if the Pf4 prophage was required for Pf virion production in biofilms. At day 6, the virion titer in biofilm effluents from the mutant were similar to those detected in the MPAO1 effluent. qPCR confirmed that these virions were exclusively derived from the Pf6 prophage (Fig. 2cd), indicating that Pf6 carriage is sufficient for virion production in biofilms. Attempts to delete the entire Pf6 prophage from MPAO1 were unsuccessful, but biofilm effluent derived from a MPAO1 derivative deleted for the Pf6 core region (MP-ΔPf6core) had a comparable virion titer as MPAO1. These virions were exclusively derived from the Pf4 prophage (Fig. 2cd), indicating that Pf4 can be produced independently of the core Pf6 prophage in biofilms. As a negative control in these experiments, no virions were detected in the biofilm effluent of MP-ΔPf4ΔPf6core (Fig. 2c), an MPAO1 derivative lacking both the Pf4 prophage and Pf6 core genes.

Pf4 production has been associated with elevated *P. aeruginosa* resistance to aminoglycoside antibiotics^15^. To examine how the presence and production of both Pf4 and Pf6 influence this phenotype, we compared the gentamicin sensitivity of biofilms derived from phage-producing strains MPAO1, MP-ΔPf4, and MP-ΔPf6core to that of the non-producer strain MP-ΔPf4ΔPf6core. Biofilms from the three phage-producing strains exhibited markedly higher resistance to gentamicin (∼30% survival) compared to the non-producer (<10^−7^% survival) (Fig. 2e), suggesting Pf4 or Pf6 production in MPAO1 biofilms promotes antibiotic resistance. In support of this idea, PAO1 biofilms, which are deficient for Pf production (Fig. 1b), and planktonic cells of all five MPAO1 strains were also highly sensitive to gentamicin. These observations suggest that virion production modifies the structure of MPAO1 biofilms in a manner that elevates resistance (or tolerance) to antibiotics. Consistent with this hypothesis, we found that mechanical disruption of biofilms derived from the three phage producing strains markedly diminished their resistance to gentamicin (Fig. 2e). Together, these data reveal that both Pf4 and Pf6 prophages give rise to Pf virions in MPAO1 biofilms, and that both Pfs modulate host cell phenotypes in biofilm conditions. Academic network searching revealed that while many groups currently use MPAO1 (often referred to as ‘PAO1’), use of Pf6-deficient PAO1 continues, suggesting that caution be taken in interpreting data involving *P. aeruginosa* strains with varying prophage carriage (Table S2 and Table S3).

## The Pf6 KKP module controls Pf lysogeny via MvaU phosphorylation

Since Pf4 production was maintained in MPAO1-ΔPf6core, but absent in PAO1 biofilms, we investigated how Pf6 controls Pf4 production. We carried out RNA-seq studies to probe the pathways that control Pf4 and Pf6 prophage activation in MPAO1 biofilms. The Pf4 and Pf6 core genes, including their respective excisionases (Pf4: *xisF4*, Pf6: *xisF6*) and major coat protein (gVIII) genes were among the most upregulated genes in MPAO1 biofilm cells versus planktonic cells, where no Pf phage particles were detected (Fig. 3a, Table S4, Extended Data Fig. 1). Conversely, no Pf4 prophage genes were significantly induced in PAO1 biofilm cells (Fig. 3a; Table S4). Thus, the presence of the two related prophages in MPAO1 appears to dramatically shift prophage gene expression in biofilms towards virion production.

**Fig. 3.**
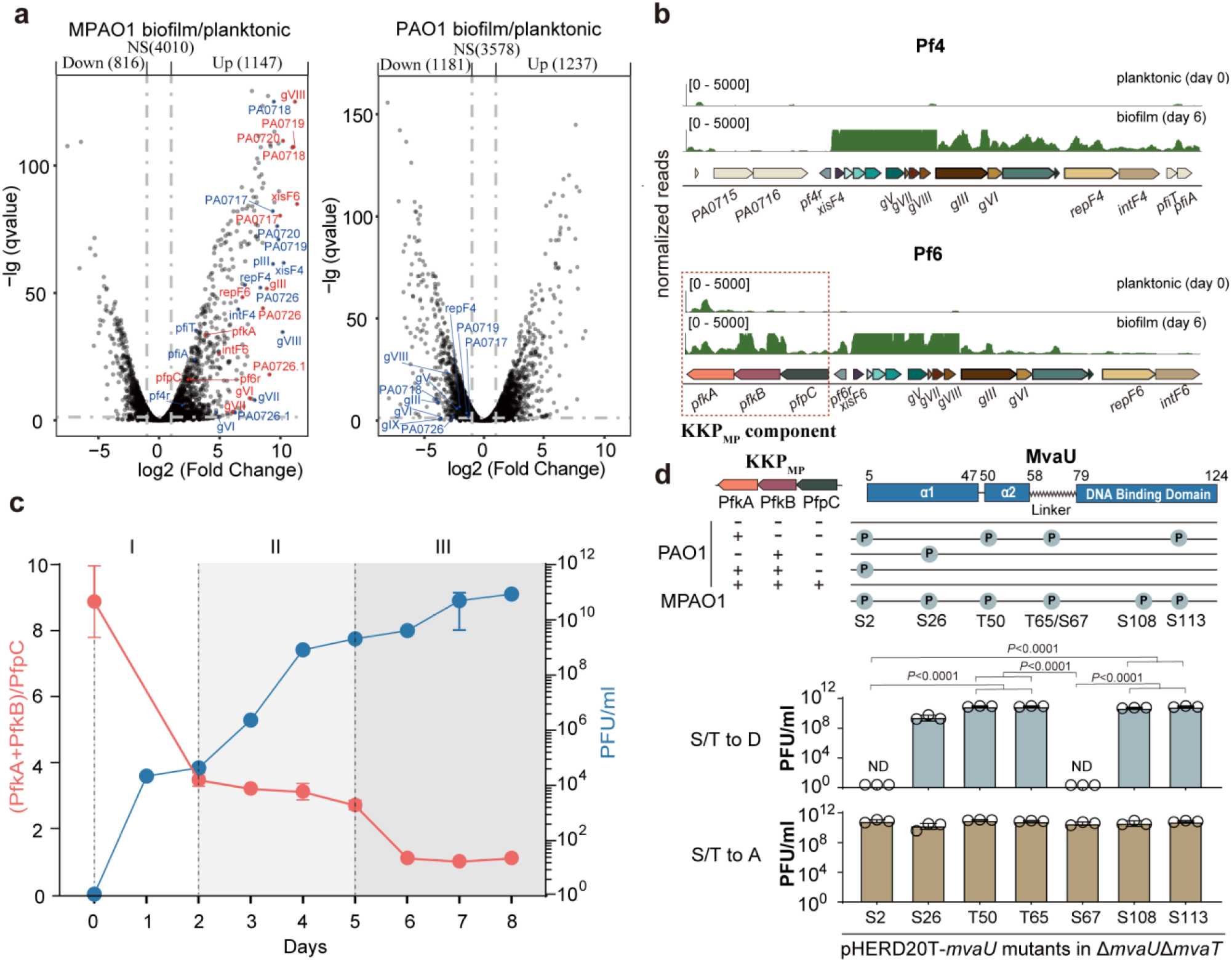
Pf6 KKP_MP_ controls Pf lysogeny via MvaU phosphorylation. **a**, Differentially-expressed genes in biofilms versus planktonic cells in MPAO1 and PAO1. Significantly changed Pf6 and Pf4 genes are labeled in red and blue, respectively. NS indicates not significantly. **b**, RNA-seq read coverage of Pf4 and Pf6 transcripts in MPAO1 biofilm and planktonic cells. Pf6 KKP_MP_ genes (*pfkA, pfkB* and *pfpC*) are highlighted. **c**, Kinetic analysis of Pf phage release and the mRNA ratio of kinases versus phosphatase (PfkA+PfkB/PfpC). Three stages are apparent: I - prophage exits lysogeny; II - phage production increases logarithmically; III - phage production plateaus. **d**, Top: Phosphorylation sites in MvaU-His purified from PAO1 or from PAO1 with indicated KKP_MP_ components introduced, or sites identified by phosphoproteomics in MPAO1 with PfkA overproduction. Bottom: Planktonic Pf phage titers from selected MvaU variants expressed in MP-Δ*mvaU*Δ*mvaT*. ND indicates not determined. Three independent cultures (*n*=3) were used for **a, c** and **d**, and data are shown as the mean ± SD in **c** and **d**.

Pf6 contains three distinctive accessory genes, two of which bear similarity to eukaryotic-like serine/threonine kinases (Pfam PF00069), and one to a serine/threonine phosphatase (Pfam PF13672). These three genes were denoted here as *pfkA, pfkB* and *pfpC*, and the entire operon as KKP_MP_ (for dual kinase phosphatase in MPAO1) (Fig. 3b, Extended Data Fig.2). We first hypothesized that KKP_MP_ enables Pf production during biofilm formation. We were unable to delete KKP_MP_ alone from MPAO1, necessitating studies of KKP_MP_ function in heterologous hosts. Introduction of the KKP_MP_ loci into the PAO1 chromosome was not sufficient to induce Pf4 production in biofilms (Extended Data Fig. 3ab), suggesting that additional factors hamper this process. Previous reports have identified a PAO1-specific N-terminal truncation in the type IV pilus component *pilC*, which is the Pf phage receptor in *P. aeruginosa*^31^ (Extended Data Fig. 3cd). Replacement of the truncated *pilC* with the full-length MPAO1 allele restored Pf biofilm production and adsorption (Extended Data Fig. 3ef), indicating an intact Pf receptor is required for high level virion production, likely by enabling superinfection. Biofilm PFUs from this strain (PAO1::*pilC*_MP_), were exceptionally high, reaching ∼ 10^10^ times the titers of MPAO1 (Extended Data Fig. 3e, Fig. 3c). Remarkably, when KKP_MP_ was transferred to PAO1:: *pilC*_MP_, Pf production was reduced to levels comparable to MPAO1 (Extended Data Fig. 3e, Fig. 3c). Thus, rather than enabling Pf production, KKP_MP_ appears to dynamically control phage production to prevent unrestrained virion release.

While *pfpC* and *pfkB* expression was primarily biofilm-restricted, *pfkA* was the highest expressed coding gene among all Pf4 and Pf6 genes in planktonic cells (Fig. 3ab, Table S4). We hypothesized that shifts in the ratio of KKP_MP_ components correlate to Pf phage production. Phage production and expression of the three KKP_MP_ components were measured daily during MPAO1 biofilm formation. Pf production in MPAO1 biofilms appeared to occur in three stages (Fig. 3c). Strikingly, the ratio of the mRNAs of the kinases versus phosphatase ((*pfkA+pfkB)*/*pfpC*) was inversely correlated to phage production (Fig. 3c). Specifically, kinase expression was at its highest in planktonic cells where Pf was not produced, and gradually decreased as Pf production rose, consistent with an inverse relationship between kinase activity and Pf prophage activation (Fig. 3c).

Since PfkA is the only KKP_MP_ component that is highly expressed when Pf gene expression is silenced, we hypothesized that a phosphorylated substrate of PfkA represses Pf gene activation. Phosphoproteomics of MPAO1 cells overexpressing PfkA identified the H-NS family protein MvaU as a target of PfkA phosphorylation (Table S5). MvaU is known to inhibit expression of the Pf4 excisionase *xisF4* and has been linked to control of Pf4 lysogeny^18^. In planktonic MPAO1 cells, MvaU was phosphorylated at multiple sites (S2, S26, T65/S67, S108, S113), but there was no detectable MvaU phosphorylation in PAO1 (Fig. 3d). Introduction of PfkA, PfkB, or both kinases into PAO1 led to MvaU phosphorylation at similar sites to those identified in MPAO1 (Fig. 3d, Extended Data Fig. 4). When we introduced PfpC along with both kinases, MvaU phosphorylation was no longer observed (Fig. 3d), highlighting that the interplay between the kinases and the phosphatase components of KKP_MP_ control the phosphorylation state of MvaU.

**Fig. 4.**
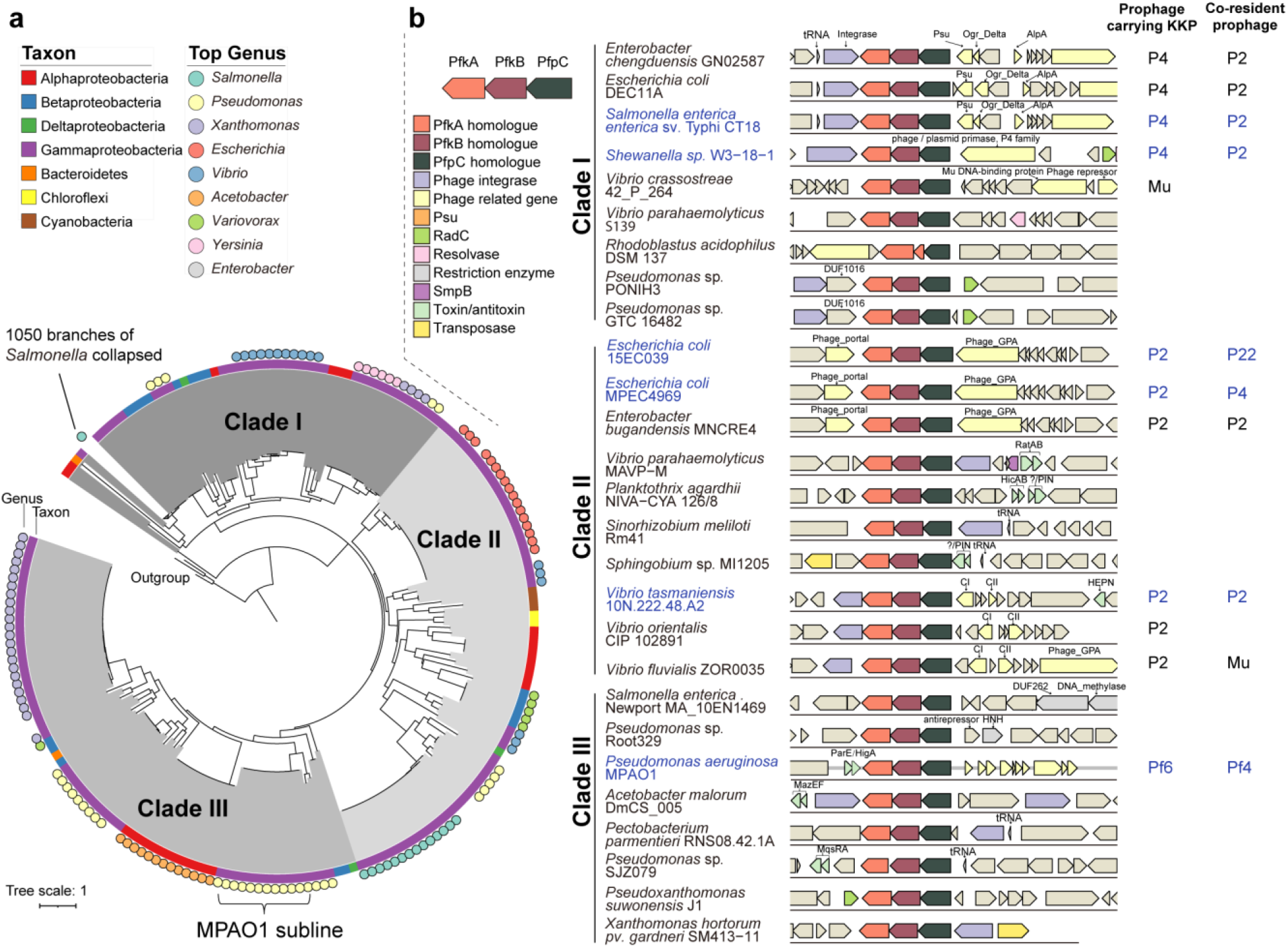
KKP modules are widespread and present in co-resident prophages. **a**, Maximum likelihood tree of PfpC-like phosphatases. Colored bars around the tree indicate taxa. Filled circles indicate genera. Many *Salmonella* sequences were collapsed. **b**, Selected KKP gene clusters and their genomic neighborhoods. KKP modules located in prophages and their co-resident prophages are indicated on the right, and other KKP-linked genes (transposons, TA systems, and restrictive modification systems) are indicated on the left. Gene names or conserved domains shown include: Psu, phage polarity suppression protein (PF07455); Phage_GPA, Bacteriophage replication gene A protein (GPA, PF05840); DNA_methylase, C-5 cytosine-specific DNA methylase (PF00145); RadC, DNA repair protein RadC (PF04002); DUF262 (PF03235); HNH, HNH endonuclease (PF01844); Ogr_Delta, viral family of phage zinc-binding transcriptional activators (PF04606); AplA, transcriptional regulator, AlpA family (PF05930); Phage_portal, Phage portal protein (PF04860); DUF1016, YhcG PDDEXK nuclease domain (PF06250); CI and CII, Bacteriophage repressor. The KKPs in prophages that were selected for follow-up are indicated in blue.

To probe whether phosphorylation of these MvaU sites influences Pf4 lysogeny, seven MvaU Ser/Thr residues in MPAO1 *mvaU* were individually substituted with either aspartate (phosphomimetic) or alanine (nonphosphorylatable) and Pf4 production was measured. These experiments were performed in an MPAO1 derivative also lacking the related H-NS family member MvaT, a deletion required for Pf virion production during planktonic growth^18,20^. Asp (D) substitutions in MvaU residues S2 or S67 totally repressed Pf4 virion production, but the corresponding Ala (A) mutations did not (Fig. 3d). Together, these data suggest a model where the high planktonic expression of *pfkA* in MPAO1 leads to MvaU phosphorylation and consequent silencing of Pf prophage gene expression. In biofilms, the elevated expression of *pfpC* leads to the dephosphorylation of MvaU, which in turn relieves repression of prophage genes that promote Pf prophage activation in MPAO1 biofilms.

## KKP gene clusters are widespread and present in co-resident prophages

Bioinformatic searches identified more than 1200 putative KKP gene clusters in Gram-negative organisms primarily in Proteobacteria, but also in Cyanobacteria and Bacteroidetes (Fig. 4a, Table S6). In most cases (1,090 instances), KKP gene clusters were located in the genomes of diverse temperate phages including P2 and its satellite P4, Mu, and Pf6-like prophages in diverse hosts (Fig. 4b). Similar to MPAO1, KKP clusters were found in several strains with multiple potentially interacting prophages. For example, *Shewanella* sp. W3-18-1 harbors a P4 satellite prophage containing a KKP cluster along with a P2-family prophage that may correspond to its helper. Similarly, *E. coli* MPEC4969 harbors a P2 prophage containing a KKP cluster along with its putative satellite P4 prophage. Thus, KKP modules may mediate interactions between co-resident prophages, as described above between Pf6 and Pf4 prophages in *P. aeruginosa*. KKP clusters that were not located in prophages were also found closely linked to genes associated with mobile genetic elements (79 out of 129), including integrases and transposons, as well as modules such as toxin-antitoxin (TA) and restriction-modification systems (Fig. 4b) that are associated with bacterial defense against mobile elements.

To examine diversity of KKP modules, we performed phylogenetic analysis of PfpC sequences and found that KKP gene clusters can be broadly classed into three clades (Fig. 4a). KKP modules of different clades were found in the same bacterial species; e.g. clade I and II modules were identified in different *E. coli* isolates, indicating that the three clades are not restricted to particular niches. Interestingly, highly similar KKP gene clusters (>90% amino acid identity) were found in *Enterobacter, Escherichia* and *Salmonella*, organisms that inhabit the gut (Fig. 4ab), suggesting that KKP gene clusters can be acquired through horizontal gene transfer.

## Prophage-encoded KKP is a tripartite TA system and provides defense against lytic phages

In addition to regulating Pf activation, we addressed whether the Pf6 KKP_MP_ module could impede infection with Pf phage. Introduction of KKP_MP_ into PAO1 provided resistance to superinfective Pf phages released from MPAO1 biofilms (Fig. 5a). Moreover, the KKP_MP_ module also conferred host protection against unrelated lytic *Pseudomonas* phages PAP8, QDWS, PAP-L5 and PAOP5 (Fig. 5ab). TA systems have recently been linked to phage defense^6,9,32-35^. The opposing functions of PfkA/PfkB and PfpC prompted us to ask if KKP_MP_ also constitutes a novel TA system. Expression of each of the three components alone in PAO1 was not toxic. However, co-expression of PfkA and PfkB drastically inhibited cell growth and viability, while co-expression of the three components together was nontoxic (Fig. 5c, Extended Data Fig. 5a). Thus, KKP_MP_ functions as a tripartite TA system where the two kinases together function as the toxin and the phosphatase as the antitoxin.

**Fig. 5.**
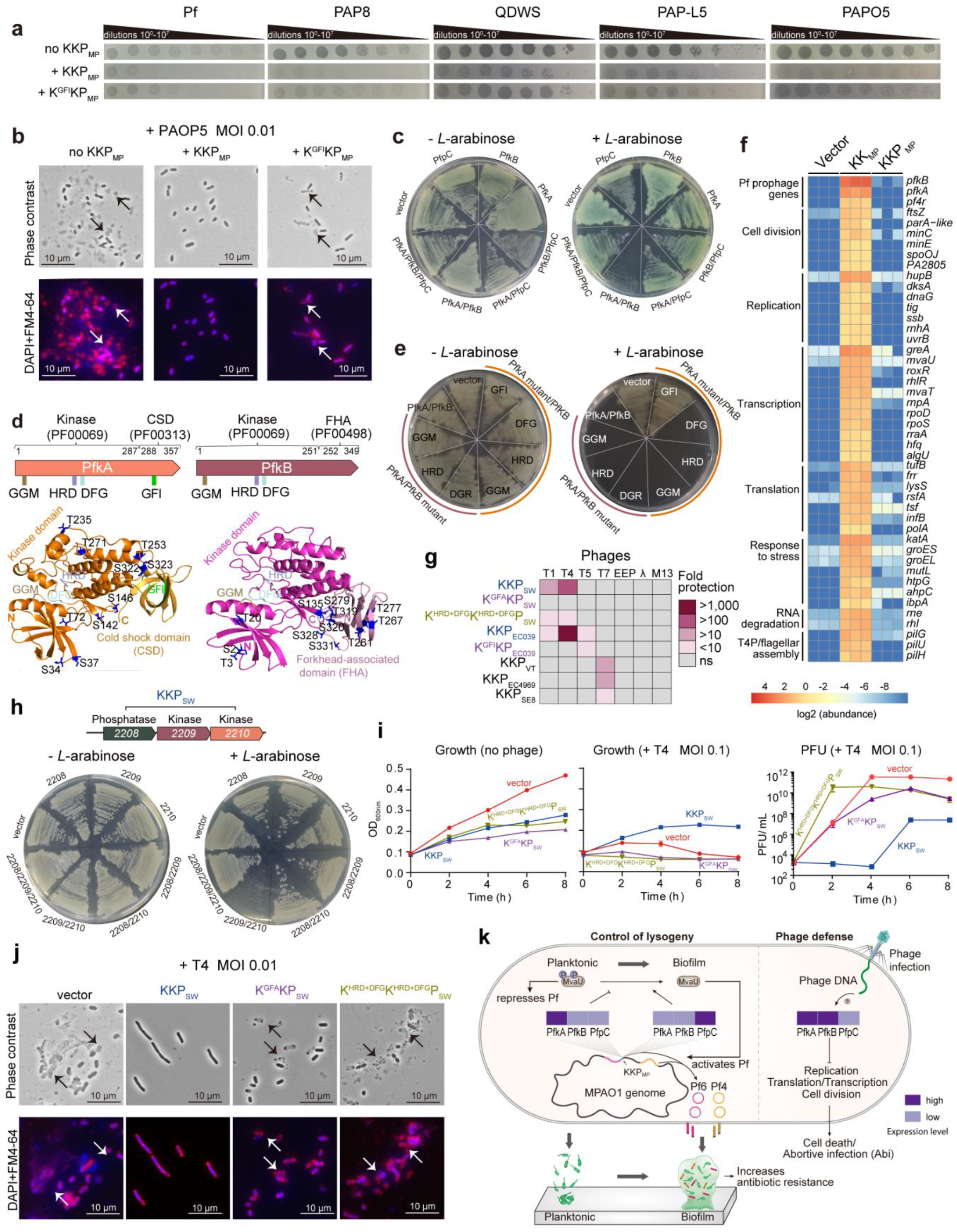
Prophage-encoded KKP is a tripartite TA and provides defense against lytic phages. **a**, Sensitivity of PAO1::*pilC*_MP_ with chromosomally integrated KKP_MP_ (+KKP_MP_) or KKP_MP_ mutant (+K^GFI^KP_MP_) to different *P. aeruginosa* phages. PAP8, PAOP5, PAP-L5 and QDWS are lytic phages. K^GFI^KP_MP_: GFI mutant of PfkA. **b**, Microscopy of FM4-64 and DAPI-stained cells (the same as **a**) after infection with PAOP5. Red, FM4-64; blue, DAPI; arrows indicate lysed cells. **c**, Toxicity of KKP_MP_ in PAO1. **d**, Conserved domains and features identified in PfkA and PfkB (top) and predicted protein structures of PfkA and PfkB (bottom). Conserved domains and phosphorylation sites identified by phosphoproteomics are labeled on the structures. **e**, Toxicity of PfkA and PfkB mutants in PAO1. GGM, HRD, DFG and GFI were changed to AAS, FEN, NYA and DYA respectively. **f**, Heat map showing a subset of the most highly phosphorylated proteins by PfkA and PfkB in PAO1. Abundance of phosphorylated proteins from PAO1 carrying pHERD20T(vector), pHERD20T-*pfkA*-*pfkB* (KK_MP_), or pHERD20T-*pfkA*-*pfkB-pfpC* (KKP_MP_) are shown using three independent cultures. **g**, Fold change protection from lysis by phages in *E. coli* K12 MG1655 expressing different KKP modules. KKP_SW_, KKP_EC039_, KKP_VT_, KKP_EC4969_ and KKP_SE8_ indicate KKP from *Shewanella* sp., *E. coli* 15EC039, *Vibrio tasmaniensis, E. coli* MPEC4969 and *S. enterica* CT8, respectively. Where indicated, superscripts denote motif(s) targeted for mutation in the corresponding expression vector. **h**, Toxicity of KKP_SW_ components. **i**, Growth curves and PFU determination for *E. coli* K12 harboring an empty vector or plasmid-encoded KKP_SW_, K^GFA^KP_SW_ and K^HRD+DFG^K^HRD+DFG^P_SW_ after infection with T4 phages at MOI 0.1. **j**, Microscopy of *E. coli* K12 cells expressing KKP_SW_, K^GFA^KP_SW_ and K^HRD+DFG^K^HRD+DFG^P_SW_ after infection with T4 at MOI of 0.01; red, FM4-64; blue, DAPI; arrows indicate lysed cells. **k**, Model for dual function of KKP_MP_ in control of lysogeny and phage defense. In planktonic cells, PfkA phosphorylates MvaU, which represses Pf activation. During biofilm development, PfpC expression is induced, leading to MvaU dephosphorylation, resulting in Pf activation. During lytic phage infection, both PfkA and PfkB are induced and phosphorylate proteins involved in essential cellular pathways, providing antiphage defence via a mechanism similar to abortive infection. Three independent cultures (*n*=3) were used in **a-c, e-j**, and data are shown as the mean ± SD.

PfkA and PfkB both have a Ser/Thr kinase domain (PF00069) with conserved GGM, HRD, and DFG motifs at their respective N-termini (Fig. 5d, Extended Data Fig. 2). The two kinases differ primarily in their C-termini. PfkA has a cold shock domain (CSD) with a conserved GFI/A motif that may have ssDNA/RNA-binding properties^36,37^. In contrast, PfkB has a Forkhead-associated (FHA) domain (PF00498) that is a phospho-specific protein-protein interaction motif in both eukaryotes and prokaryotes^38^ (Fig. 5d). To examine the role of these sequences in PfkA and PfkB toxicity, we generated a panel of motif mutants and tested their toxicity in PAO1. Single or multiple-motif mutations in PfkA or PfkB alone did not alleviate their toxicity (Fig. 5e; Extended Data Fig. 5b). Only double-motif mutations of HRD or DFG in both PfkA and PfkB abolished toxicity (Extended Data Fig. 5b), suggesting the two kinase domains have some functional complementarity. Point mutations in the GFI motif of the CSD domain of PfkA alone also completely abolished toxicity (Fig. 5e). Notably, truncations revealed that intact kinase domains are not sufficient for exerting the kinases’ toxicity (Extended Data Fig. 5c), suggesting that parts of the C-terminal domains of PfkA and PfkB are critical for their function.

Since KKP_MP_ requires two Ser/Thr kinases to exert toxicity, we used phosphoproteomics to identify host targets that are phosphorylated in the presence of PfkA and PfkB. We identified many probable PfkA/B targets that are involved in essential cellular pathways, including proteins in cell division (*ftsZ, minC, minE*), DNA replication (*dnaG, dksA*), transcription (*mvaU, mvaT, rpoS, rpoD*), and translation as well as stress-related genes. Expression of PfpC alongside PfkA and PfkB returned phosphorylation to nearly background levels (Fig. 5f, Table S7). We also observed high levels of PfkA and PfkB phosphorylation in the absence of PfpC, suggesting PfkA/B are activated by phosphorylation and deactivated by PfpC activity. Toxin neutralization via dephosphorylation suggests KKP_MP_ represents a type VII TA system, in which the antitoxin (PfpC) inactivates the toxin (PfkA/B) by post-translational chemical modification^39,40^. Importantly, the PfkA GFI mutant conferred reduced resistance to phage infection in PAO1 (Fig. 5ab), suggesting that KKP_MP_ may provide phage defense via kinase toxicity leading to abortive infection (Fig. 5bde, Extended Data Fig. 5bc).

Finally, to examine the scope and conservation of KKP anti-phage activity, we tested KKP modules derived from five unrelated and non-filamentous prophages of *Shewanella* sp. W3-18-1, *Vibrio tasmaniensis, E. coli* 15EC039, *E. coli* MPEC4969 and *Salmonella enterica enterica* sv. Typhi str. CT18 in a heterologous *E. coli* K12 expression system. These KKP modules each conferred protection against at least one canonical lytic phage, including T1, T4 or T7 (Fig. 5g; Extended Data Fig. 6 and 7). Further characterization of KKP_SW_ from *Shewanella* sp. W3-18-1, which is part of a P4 prophage, confirmed that it is a tripartite TA system akin to the KKP_MP_ (Fig. 5h; Extended Data Fig. 6a). The anti-phage activity of the *Shewanella* and *E. coli* 15EC039 KKP modules was abolished when the conserved GFA or GFI motif in their respective PfkAs were mutated. Furthermore, mutations of HRD and DFG motifs in both PfkA and PfkB of *Shewanella* also markedly reduced anti-phage activity, suggesting that the mechanism of KKP-mediated phage defense is conserved (Fig. 5ij; Extended Data Fig. 6b and 7). Thus, prophage-encoded KKP modules can defend host cells against infection by diverse lytic phages.

## DISCUSSION

Provision of both prophage regulatory and phage defense functions by the three gene KKP cassette unites two functionally disparate activities in an unprecedented fashion. The widespread conservation of KKP modules in Gram-negatives suggests that this phosphorylation-based control system is a common mechanism of host-phage and host-prophage crosstalk. Nearly 90% of the >1200 identified KKP modules were in prophages, suggesting that diverse phyla have a dual-functional KKP. However, given the identification of >100 KKP cassettes outside of prophages, often within other mobile elements, it is possible some KKP clusters only mediate phage defense. In addition, this regulatory module is often present in microbes that have co-resident prophages and, based on findings with the co-resident Pf6 and Pf4 in *P. aeruginosa*, we propose that KKP modules also mediate interactions between prophages in other organisms. The functions of KKP in any given bacterium are likely shaped by prophage content, the frequency and type of lytic phage attack, as well as host-intrinsic phage control mechanisms.

Accessory genes in prophages, e.g., prophage-borne toxins like the cholera toxin genes in CTXΦ^11^, often bestow critical properties to their hosts, but usually do not directly modify prophage development as we found with KKP_MP_. Phosphorylation of a host H-NS protein (MvaU) by KKP_MP_’s dual kinases, PfkA and PfkB, represses expression of Pf genes required for prophage activation, keeping Pf6 and Pf4 gene expression largely silent in planktonic cells. However, in biofilms, expression of the PfpC phosphatase alleviates Pf gene repression, presumably via dephosphorylation of MvaU, inducing production of Pf4 and Pf6 virions, which alter the properties of *P. aeruginosa* biofilms, elevating antibiotic resistance and virulence (Figure 5k)^5,15,17^. Prophage induction is often regulated by control of phage repressor activity, e.g. repressor cleavage by host factors^41^, but reversible kinase-phosphatase control of lysogeny by prophage-encoded factors has not been described. This mechanism enables dynamic prophage-host crosstalk and places KKP at the intersection of environmental signals, prophage activation, and host physiological state.

KKP is a phosphorylation-based tripartite TA system, where the PfkA and PfkB kinases serve as the toxin and the PfpC phosphatase as the antitoxin. KKP’s phage defense function relies on the toxic properties of its kinases, which target many essential cell processes. Several recent studies have uncovered that both phage- and chromosome-encoded TA systems mediate phage defense by leading to abortive infection (Abi)^6,32-35^ and our findings suggest that KKP promotes defense through a similar process. Since KKP functions through post-translational modifications that are potentially reversible, in contrast to other TA-based mechanisms of phage defense, this module could provide enhanced host control of cell fate. Identifying the pathways that relay diverse signals (e.g., phage attack or biofilm formation) to ultimately regulate KKP output may reveal novel triggers of phage defense and prophage induction. The discovery of KKP expands understanding of how mobile genetic elements such as phages can become entwined with microbial host pathways and modulate host phenotypes. The diversity of bacteria-phage interactions has become increasingly apparent with the advent of high-throughput screening and enrichment analyses^35,42-46^. However, none of the known phage defense systems identified to this point involve a reversible phosphorylation cascade orchestrated by complementary kinases and a phosphatase, suggesting that KKPs represent a new axis through which bacteria can respond to phage attack. Given that several other newly identified phage defense systems are prophage-encoded^6-9^, it will be of interest to investigate whether these systems also control additional types of phage-host or phage-phage interactions, such as prophage activation like KKP, or the establishment of lysogeny.

## ACKNOWLEDGMENTS

This work was supported by the National Science Foundation of China (91951203, 31970037, 31625001 and 42188102), by the Guangdong Major Project of Basic and Applied Basic Research (2019B030302004), the Guangdong Local Innovation Team Program (2019BT02Y262), the Innovation Academy of South China Sea Ecology and Environmental Engineering, the Chinese Academy of Sciences (ISEE2018PY01), and the Key Special Project for Introduced Talents Team of Southern Marine Science and Engineering Guangdong Laboratory (Guangzhou) (GML2019ZD0407). The Waldor lab is supported by HHMI. This article is subject to HHMI’s Open Access to Publications policy. HHMI lab heads have previously granted a nonexclusive CC BY 4.0 license to the public and a sublicensable license to HHMI in their research articles. Pursuant to those licenses, the author-accepted manuscript of this article can be made freely available under a CC BY 4.0 license immediately upon publication.

## AUTHOR CONTRIBUTIONS

X.X.W., M.K.W. and Y.X.G. conceived the project. Y.X.G., K.H.T. and B.S. contributed equally to this work. Y.X.G., J.Y.G., J.Z.L., S.T.L., X.X.L., W.Q.W., X.Y.G., Z.L.N. and T.L.L. constructed strains, performed all other wet-lab experiments, K.H.T., Y.X.G., B.S. and R.C. conducted all bioinformatic analyses. All authors interpreted data, X.X.W., M.K.W., B.S., Y.X.G. and K.H.T. wrote the paper.

## ADDITIONAL INFORMATION

Supplementary information is available for this paper. Correspondence and requests for materials should be addressed to XXW.

## METHODS

### Bacterial strains, plasmids and growth conditions

The bacterial strains, plasmids and all primers used in the study are listed in Table S8 and Table S9. *P. aeruginosa* PAO1 and MPAO1 sublines and *Escherichia coli* were cultured at 37°C at 220 rpm in Luria-Bertani (LB) medium. Gentamicin (Gm, 30 µg ml^−1^) and ampicillin (50 µg ml^−1^) were used to maintain pEX18Gm-based and pFLP2 or pEX18Ap-based and pHERD20T-based plasmids, respectively.

### Flow-cell biofilm assay

Flow-cell biofilm assays were performed as previously described with minor modifications^47^.The system was assembled using sterilized material as shown in Fig. 1a. The assay was performed at room temperature with modified M9 medium containing 47.8 mM Na_2_HPO_4_, 22 mM KH_2_PO_4_, 6.8 mM NH_4_Cl, 18.7 mM NaCl, 100 µM CaCl_2_, 2 mM MgSO_4_ and 0.1% glucose. For inoculation, 1 ml fresh overnight cultures were injected into the inlet of medical silicone catheters (inner diameter × length, 3 mm × 400 mm) (Forbest Manufacturing Corporation, Shenzhen, China), taking care to avoid any bubbles. Inoculated catheters were left static for 1 hour before flow initiation to allow colonization on the inner catheter surface. Then, flow was initiated with a peristaltic pump with a flow rate of 0.1 ml min^−1^ for each channel to wash away unattached cells. At indicated time points, phages were collected from the effluents and biofilm cells inside the catheter were collected by cutting the catheter open. Fresh M9 medium was supplied daily and total assay runtime lasted for 6-9 days depending on the readout.

### Quantification of Pf phages

Phage plaque assays were conducted using the top-layer agar method^48^ to quantify the number of Pf phages. Briefly, 2 ml of biofilm effluents were collected at day 6 post-inoculation and centrifuged at 13,000 × g for 5 mins. The obtained supernatants were filter sterilized with 0.22 μm filters (Millipore Corporation, Villerica, MA, USA) to obtain cell-free phage solutions. The solutions were serially diluted by 10-fold and 5 μl of each dilution was spotted on a R-top layer medium contained MPAO1 or PAO1 wild type or deletion strains. The cell-containing layer was prepared by mixing 500 μl of fresh overnight cultures with 5 ml of 55°C R-top medium containing 0.8% (w/v) agar, 0.1% yeast, 1% tryptone, and 1% NaCl. The R-bottom layer contains 1% (w/v) agar, 0.1% yeast, 1% tryptone and 0.8% NaCl. After drying, plates were cultured at 37°C for 16 h and the numbers of plaques formed were counted. For plaque assays involving overexpression of genes, overnight cells were 1:1000 diluted and cultured to turbidity (OD_600_ ∼ 1). 10 mM *L*-arabinose was then added for 2 h to induce gene expression, after which cells were mixed with R-top medium containing 10 mM *L*-arabinose and plaque assays performed as above.

### Quantitative PCR (QPCR)

For Pf phage, phages in effluents collected at day 6 were 10-fold serial-diluted and the levels of Pf4-specific gene *pfiA* and Pf6-specific gene *pfkA* were determined using standard qPCR methods^49^. The linear regression between Ct values and phage titers for each primer set was used to determine the gene copies (gc) of *pfiA* and *pfkA* in effluents collected each day. For bacterial cells, genomic DNA of strains and related Pf prophage mutation strains were isolated using the TIANamp Bacteria DNA kit (Tiangen Biotech Co. Ltd, Beijing, China) according to the manufacturer’s instructions. During the isolation process, RNA was removed by adding 10 U/µl of RNase A during cell lysis process. A total of 20 ng DNA of each sample was used to quantify copies of the above two Pf genes. The *gyrB* was used to normalize the copies of both Pf genes in bacterial cells. Standard curves for primers used in this study were determined with serial dilutions of MPAO1 genomic DNA, and all the primers showed well amplification efficiencies.

### Microscopy

For transmission electron microscopy (TEM), phages in filtered effluents at day 6 were negative stained with 2% phosphotungstic acid. Phage ultrastructure was then imaged with a TEM H-7650 (Hitachi) instrument at the Guangdong Institute of Microbiology (Guangzhou, China). For fluorescence microscopy of GFP-tagged MCP (pVIII), biofilms formed in medical catheter at day 6 were collected and observed with a Scope.A1 fluorescence microscope (Carl Zeiss, Jena, Germany). For fluorescence microscopy during anti-phage assays, overnight cells were diluted and grown in medium with 10 mM *L*-arabinose induction at 37°C till OD_600_ ∼ 1. Phages were then added to cultures at the indicated MOIs for 3 h. Cells were then stained with FM4-64 (0.5 µg ml^−1^) and DAPI (1 µg ml^−1^) for 10 min and subsequently imaged.

### Chromosomal gene knockout and insertion in MPAO1/PAO1

A previously described gene knockout method was used to construct chromosomal gene deletion and insertions in *P. aeruginosa* ^50^ except for those we have constructed previously^18^. All primers used are listed in Table S9. For new deletion strains, the upstream and downstream fragments of each gene were PCR-amplified from MPAO1/PAO1 genomic DNA. Gel-purified amplicons were then ligated into the modified suicide plasmid pEX18Gm. The constructs were confirmed by sequencing using primers pEX18Gm-F/R. Then, deletion vectors were transferred into *E. coli* WM3064, a diaminopimelic acid (DAP) auxotroph, and conjugated into MPAO1/PAO1 strains. In-frame deletion mutants were obtained via homologous recombination using the sucrose resistance selection method. For constructions that were not successful with this method, the Gm resistance gene cassette from the plasmid pPS856 was PCR-amplified and inserted between the gene-up and gene-down fragments to enable antibiotic selection of recombinants. The Gm resistance cassette was removed from the chromosome using plasmid pFLP2 as described previously in final strains^50^. Final deletion mutants were confirmed by PCR using primers gene-SF/SR, gene-LF/LR and DNA sequencing. For chromosomal insertion constructs, the KKP_MP_ or K^GFI^KP_MP_ mutant with their own promoters were PCR fused with the Gm resistance gene, then the obtained products were fused with gene upstream and downstream fragments and ligated into pEX18Ap.

### Antibiotic resistance assay

To assess the antibiotic resistance of planktonic cells, exponentially growing cells (OD_600_ ∼1) were treated with 20 µg ml^−1^ gentamicin (Gm) for 1 h, and colony formation units (CFU) were determined by serial plating. To assess the antibiotic resistance of biofilm cells, attached biofilms were removed from inside catheters by cutting them open; the biofilms were then either mechanically disrupted by vortexing vigorously or left intact. CFU were determined after 3 h treatment with Gm.

### Construction of plasmids and site directed mutagenesis

For the construction of all the expression plasmids, the PCR amplified fragments were ligated into pHERD20T plasmid digested with NcoI and HindIII as described previously^18^. For the cloning of genes with specific site mutagenesis, previously reported procedures were used^51^.

### *de novo* sequencing and assembling of PAO1 and MPAO1 genomes

For genome sequencing of PAO1 and MPAO1, genomic DNA were isolated from exponential phase (OD_600_ ∼ 1) cells. gDNA was further purified using Agencourt AMPure XP Kit (Beckman Coulter). Then the purity, quantity, and size of DNA was assessed through a combination of NanoDrop, Qubit and pulsed field gel electrophoresis. Following quality control, 5 μg DNA was used to prepare 20-kb SMRTbell libraries per the manufacturer’s directions (Tianjin Biochip Corporation, Tianjin, China). Subsequently, the quantity and size of each library was assessed and sequenced on a Pacific Biosciences (PacBio) RSII instrument using P6-C4 chemistry with 6 h movie time following the manufacturer’s recommendations. A total of 108,077 reads for PAO1 and 32,240 reads for MPAO1 were obtained, and they were assembled using HGAP algorithm version 4 into one contig with average genome coverage of 800 × and 300 × respectively. The genome sequences of MPAO1 and PAO1 were compared with the first sequenced PAO1 genome (Genebank ID: AE004091) and MPAO1 (Genebank ID: PRJNA438597). The genome sequences of MPAO1 in this study was used as a reference. Images showing the four genome comparisons were generated using BLAST Ring Image Generator (BRIG) ([version 0.95-dev.0004](http://brig.sourceforge.net/brig-development-version-0-95-dev-0004-has-been-released/))^52^.

### Phage adsorption assay

The phage adsorption assay we performed as we reported recently with minor modification^53^. Briefly, 2 ml of culture was collected at OD_600_ ∼ 1 and mixed with different amounts of phages (10^7^ -10^9^ PFU/ml) to reach a MOI of 0.01. Then the phage and cell mixtures were cultured at 37°C for 10 min to allow phages to adsorb to the cells, and supernatants were plated to quantify PFUs before and after adsorption.

### Quantitative real-time reverse-transcription PCR (qRT-PCR)

Total RNA in planktonic and biofilm cells of PAO1 and MPAO1 were isolated using the Bacteria RNAprep pure kit (Tiangen Biotech Co. Ltd, Beijing, China) according to the manufacturer’s instructions. During the isolation process, DNA was removed by adding 10 µl of DNase I (10 U/µl) and incubating for 30 min at 37 °C. A total of 200 ng total RNA was used for reverse transcription reactions to synthesize cDNA with revere transcription system A3500 (Promega, Madison, WI). Then, 50 ng of cDNA was used to conduct qRT-PCR with SYBR green reaction mix on a Step One Real-Time PCR System (Applied Biosystems). 16S rRNA gene levels were used to normalize expression levels of KKP components in MPAO1 biofilms at different days. The fold changes of genes were calculated based on expression level of *pfkB* at day 0.

### Strand-specific RNA sequencing

Overnight PAO1 and MPAO1 cultures in M9 medium (OD_600_ ∼3) were collected by centrifugation at 12,000 g for 30 s at 4°C. The biofilm cells on the inner surface of the medical catheter were collected at day 6 by cutting 5 mm at the middle of the catheter, and cells were collected by centrifugation. Total RNA was isolated and quality control was performed with Agilent RNA ScreenTape (Lot: 0201937-152). The rRNA in the sample was removed by hybridization with the rRNA probe using Ribo-off rRNA Depletion Kit (Bacteria) (Vazyme, # N407-C1). To prepare cDNA libraries, 1µg total RNA was used for cDNA synthesis amplification (11 cycles) using VAHTS Total RNA-seq (Bacteria) Library Prep Kit for Illumina (Lot:7E582A1) according to the manufacturer’s instructions. Then, the libraries were sequenced on a NovaSeq 6000 S4 (Illumina). The obtained raw data were evaluated with FastQC (v0.10.1), and contamination and adapter sequences were removed with Cutadapt (version 1.9.1). Cleaned data were mapped to the MPAO1/PAO1 genome with bowtie2 (v2.2.6), and the expression profiling was calculated with Htseq (V 0.6.1). The differential gene expression (DGE) analysis was performed using EdgeR (R-3.1.2), and significantly changed genes (≥2-fold and FDR≤0.05) in biofilm compared to planktonic cells were selected for further KEGG analysis.

### Phosphoproteomic analysis

For PfkA overproduction, the MPAO1 cells harboring pHERD20T-*pfkA* and empty vector pHERD20T were grown to OD_600_ ∼4 in LB medium with 0.3% *L*-arabinose and collected by centrifugation at 6,000 g for 5 mins at 4°C. The cell pellets were washed twice with TBS and stored at -80°C. For KK_MP_ and KKP_MP_ overproduction, the PAO1 cells harboring pHERD20T, pHERD20T-*pfkA*-*pfkB* and pHERD20T-*pfkA*-*pfkB-pfpC* were allowed to grow for 4 h and then expression of genes was induced with 0.3% L-arabinose for 1.5 h and then cells were sampled as described above. Lysate preparation and LC-MS/MS analysis were carried out by Jingjie PTM BioLab (Hangzhou, China) Co. Ltd. For preparation of protein lysate, samples were sonicated at 220 W for 25 times (3 s per time, 5 s interval) on ice by XM-900T sonicator (Ningbo Xinzhi Biological Technology Co. Ltd, Ningbo, China) in lysis buffer (8 M urea, 1% protease inhibitor cocktail, 1% phosphatase inhibitor). The remaining debris was removed by centrifugation at 12,000 g at 4 °C for 10 min. Finally, the supernatant was collected and the protein concentration was determined with the bicinchoninic acid assay according to the manufacturer’s instructions. The procedures of protein trypsinization, phosphopeptide enrichment, LC-MS/MS, and data analysis were conducted as previously described^54,55^. The resulting MS/MS data were processed using MaxQuant search engine (v.1.6.15.0), and the tandem mass spectra were searched against the MPAO1 coding genome.

### Protein purification and LC-MS/MS analysis

MvaU with six histidines at the C-terminus was purified in PAO1 and MPAO1 cells expressing pHERD20T-*mvaU-*His, pHERD20T-*mvaU*-His-*pfkA*, pHERD20T-*mvaU*-His-*pfkB*, pHERD20T-*mvaU*-His-*pfkA-pfkB* or pHERD20T-*mvaU*-His-KKP. All the strains were cultured in LB with carbenicillin and induced with 0.3% *L*-arabinose for 3 h. Cells were collected and His-tagged proteins purified as described previously^51^. Phosphorylation sites of the purified MvaU-His in different hosts as indicated in Fig. 3d were identified as previously described^55^.

### Phylogenetic analysis of KKP modules

Gene clusters with dual kinases and phosphatase in prokaryotic genomes were identified using the Function Profile analysis tool in the IMG/M database^56^ with the following functions: phosphatase: COG0631 or PF13672, kinase: COG0515 or PF00069. All retrieved genes were further filtered by gene orientation and gene length (<1300 bp and >900 bp).The filtered sequences of phosphatases were aligned by MAFFT^57^ and further edited by trimAl^58^. Each final data set was used for the maximum likelihood (ML) phylogenetic analysis by the W-IQ-TREE^59^. The best-fit substitution model was automatically determined and the reliability of internal branches was tested by 1000 ultrafast bootstrap replicates^60^ in the W-IQ-TREE web interface. The tree was further annotated by the iTOL tool^61^.

### Protein 3D structure prediction

Protein structures of PfkA and PfkB were predicted with RobeTTAFold^62^, and Dali server^63^ was used to search for structural homology. The structural figures were produced with PyMOL (www.pymol.org).

### Growth curves after phage infection

Overnight *E. coli* cells harboring the indicated KKP and mutant plasmids were diluted to OD_600_ of 0.2. 100 µl of culture was seeded into 96-well plates, then 100 µl of indicated phages with different phage titers was mixed with seeded cells to obtain the indicated MOIs. The mixtures were allowed to grow at 37°C without shaking, and OD_600_ was measured at the indicated timepoints.

### One-step growth curves for measuring anti-phage activity

The experiment was conducted as described recently^6^. Specifically, cells with the indicated plasmids were infected with phages at MOI in LB medium containing ampicillin and *L-*arabinose, then the cells were cultured at 37°C at 220 rpm. Phages were collected at indicated time points by centrifugation at 2,400 g for 5 min and sterile-filtering the supernatant with 0.22 µm filters. The obtained phages were serially diluted and immediately dropped onto top medium containing 0.5% agar and MG1655 *E. coli* cells to quantify the phages.

### Statistical analysis

For all the experiments, at least three independent biological replicates were used. Significance testing was performed by an unpaired t-test for comparisons between two groups, and a one-way analysis of variance (ANOVA) test with Tukey’s correction for multiple comparisons. A probability level (*p*-value) of <0.05 was considered as statistically significant.

### Data availability

The data that support the findings of this study are provided within the manuscript and its associated Supplementary Information. Bacterial genome sequences of MPAO1 and PAO1 are deposited in NCBI’s GeneBank Database under BioProject PRJNA745538, and the accession numbers are CP079712 and CP085082. They were annotated by combination of the annotation to MPAO1(Genebank accession No. PRJNA438597) and RASTA rapid annotation using subsystem technology (version 2.0)^64^. In addition, the draft genome of *E. coli* 15EC039 has been submitted to the same database under accession number JAMFTL000000000. RNA-seq raw data are deposited in Sequence Read Archive (SRA) under BioProject accession number PRJNA835970. The mass spectrometry data of MvaU purified from different hosts, the mass spectrometry phosphoproteomics data with PfkA overproduction in MPAO1 and the mass spectrometry phosphoproteomics data with KK and KKP overproduction in PAO1 have been deposited to the ProteomeXchange Consortium via the PRIDE^65^ partner repository with the dataset identifier PXD035222, PXD035424 and PXD035228 respectively. There are no restrictions on data availability. Source data are provided with this paper.

## EXTENDED DATA FIGURES AND LEGENDS

**Extended Data Fig. 1.**
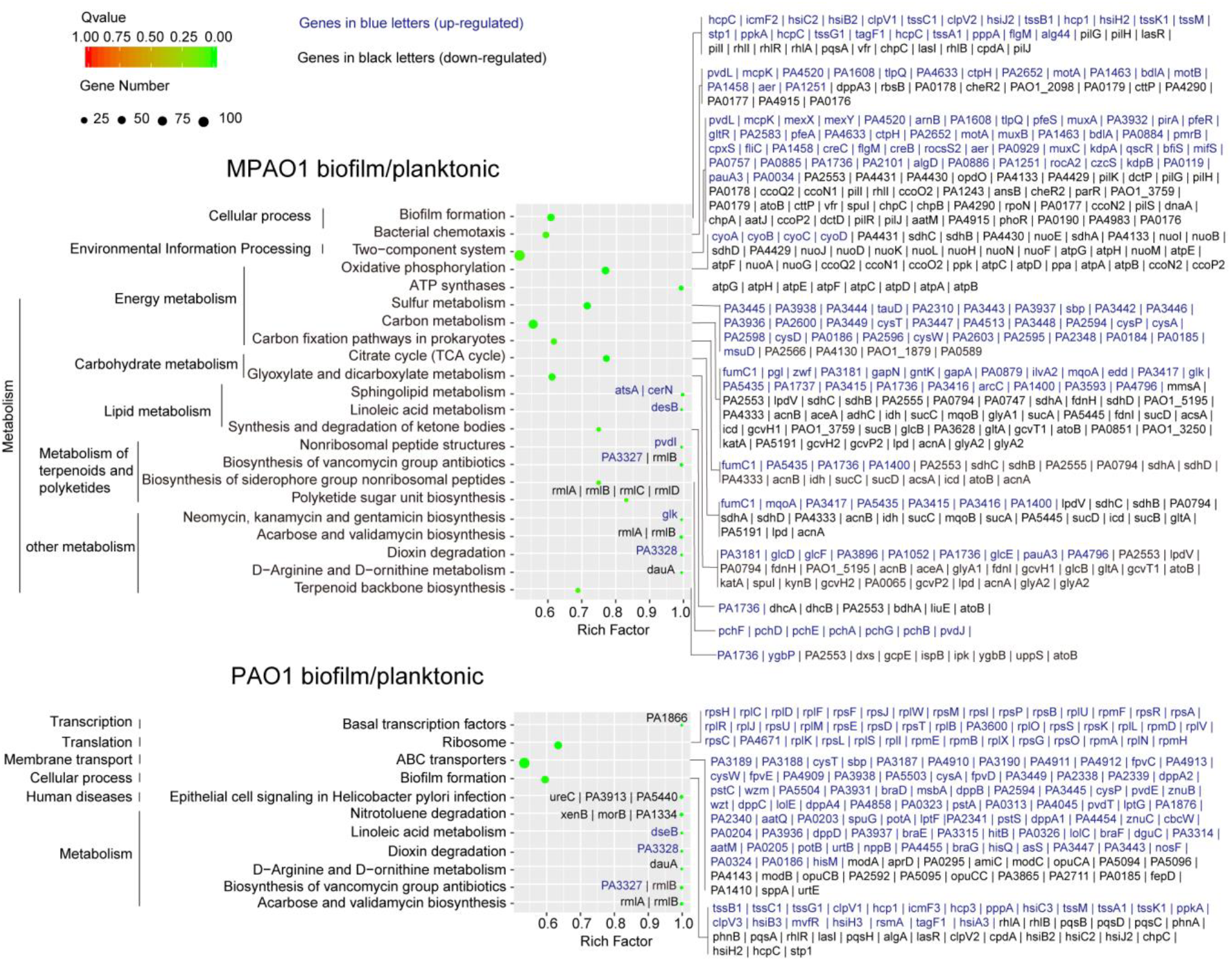
Pathway enrichment analysis of MPAO1 and PAO1 biofilm and planktonic transcriptomes. Upregulated genes are shown in blue and downregulated genes are shown in black. Number of genes enriched in each gene set is shown by the circle size. KEGG pathways with Q value <0.05 were selected as significant.

**Extended Data Fig. 2.**
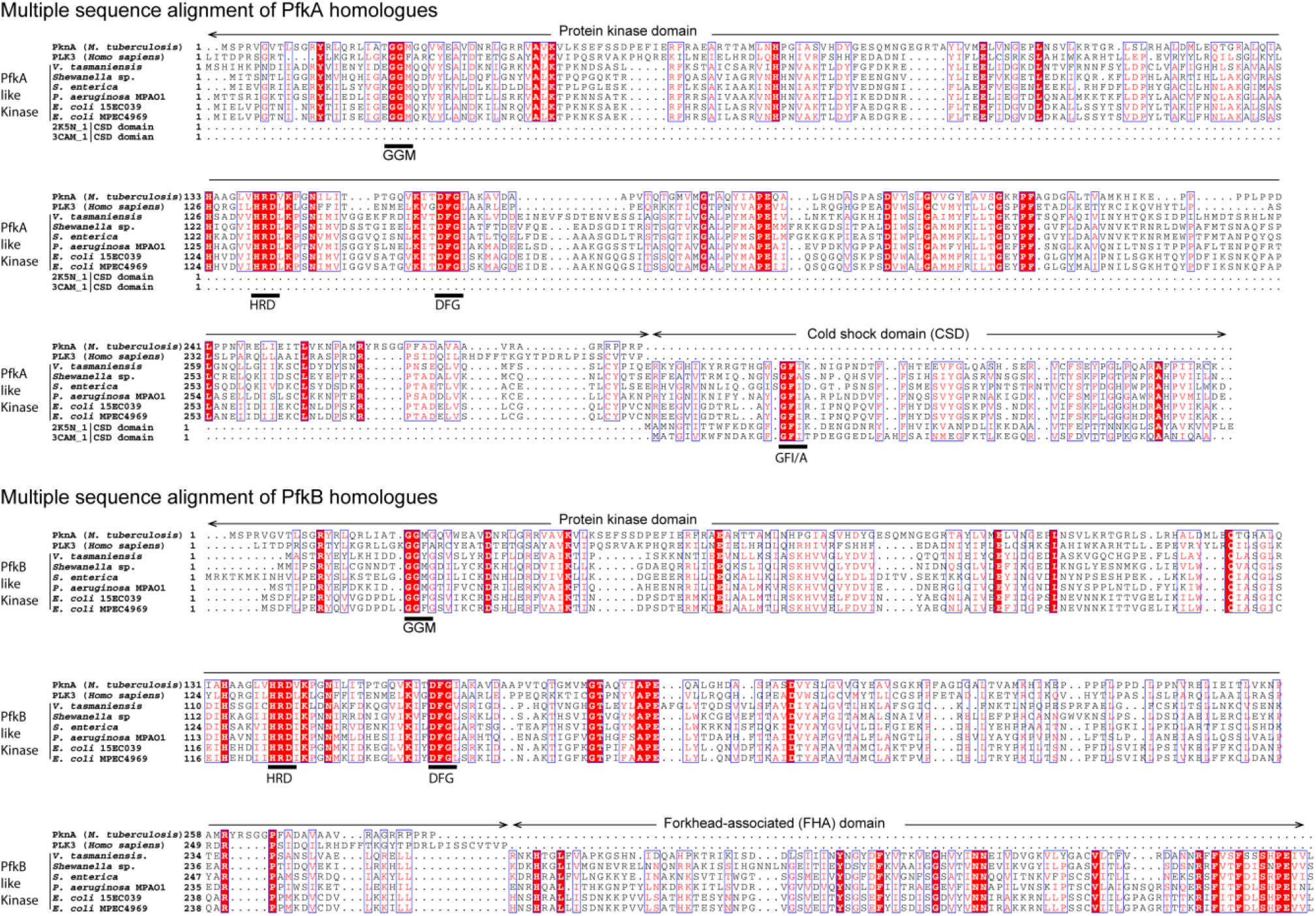
Sequence alignment of PfkA and PfkB homologues. Homologues of PfkA and PfkB used in this study were aligned with Ser/Thr kinase PknA of *Mycobacterium tuberculosis* (NCBI accession number NP_214529) and Ser/Thr kinase PLK3 of *Homo sapiens* (NCBI accession number NP_004064). Accession numbers of PfkA and PfkB homologues from *Vibrio tasmaniensis* 10N.222.48. A2, *Shewanella* sp. W3-18-1, *Salmonella enterica* enterica sv. Typhi CT18 and *E. coli* strains are listed in Table S6. The conserved domains mutated in Fig. 5 and Extended Data Fig. 5, 6 and 7 are underlined.

**Extended Data Fig. 3.**
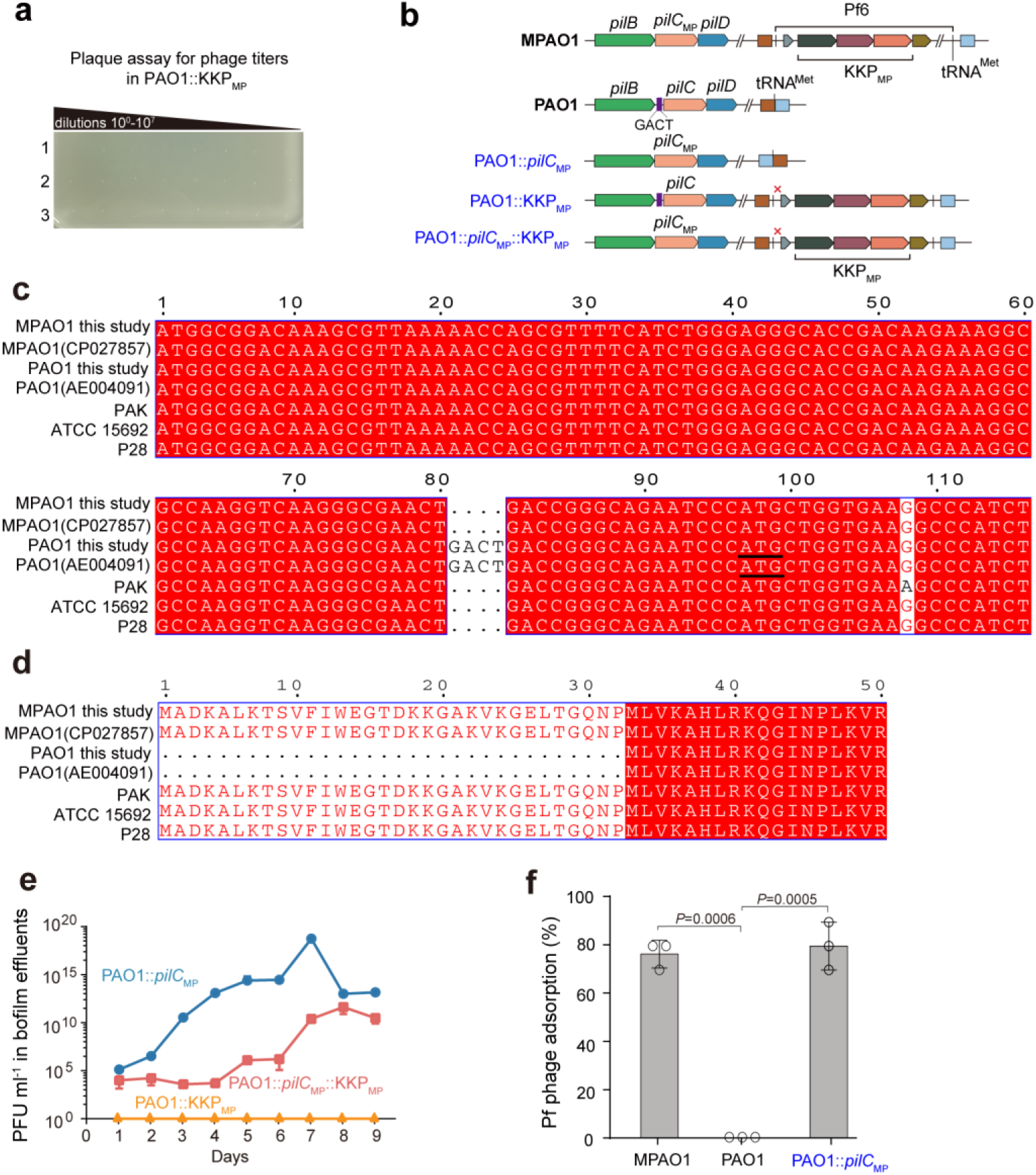
Pf phage release is controlled by KKP_MP_ in PAO1::*pilC*_MP_ biofilms. **a**, Effect of introducing KKP_MP_ in PAO1 on phage release during biofilm formation. Each row represents a biological replicate at day 9. **b**, Schematic of generated mutants. PAO1 has a 4-bp insertion in *pilC* which generates an alternative start codon and results in a loss-of-function 32 aa truncation at N-terminal PilC. The four mutants with chromosomal modification in either PAO1 or MPAO1 are shown in blue. To ensure Pf6r was not expressed in the KKP_MP_ cassette, the start codon of *pf6r* was mutated to a stop codon (marked with a red cross). **c**, Multiple alignment of *pilC* genes (first 116 bp of *pilC*_MP_) in different *P. aeruginosa* strains, and the start codon of truncated *pilC* in the two PAO1 strains are underlined. **d**, Multiple alignment of the N-terminal 50 aa of PilC in different *P. aeruginosa* strains. **e**, Pf phage titers in biofilm effluents from PAO1::KKP_MP_, PAO1::*pilC*_MP_ and PAO1::*pilC*_MP_::KKP_MP_. **f**, Adsorption of Pf phages to MPAO1, PAO1 and PAO1::*pilC*_MP_. Three independent cultures were used in **a, e**, and **f**, and data are shown as the mean ± SD.

**Extended Data Fig. 4.**
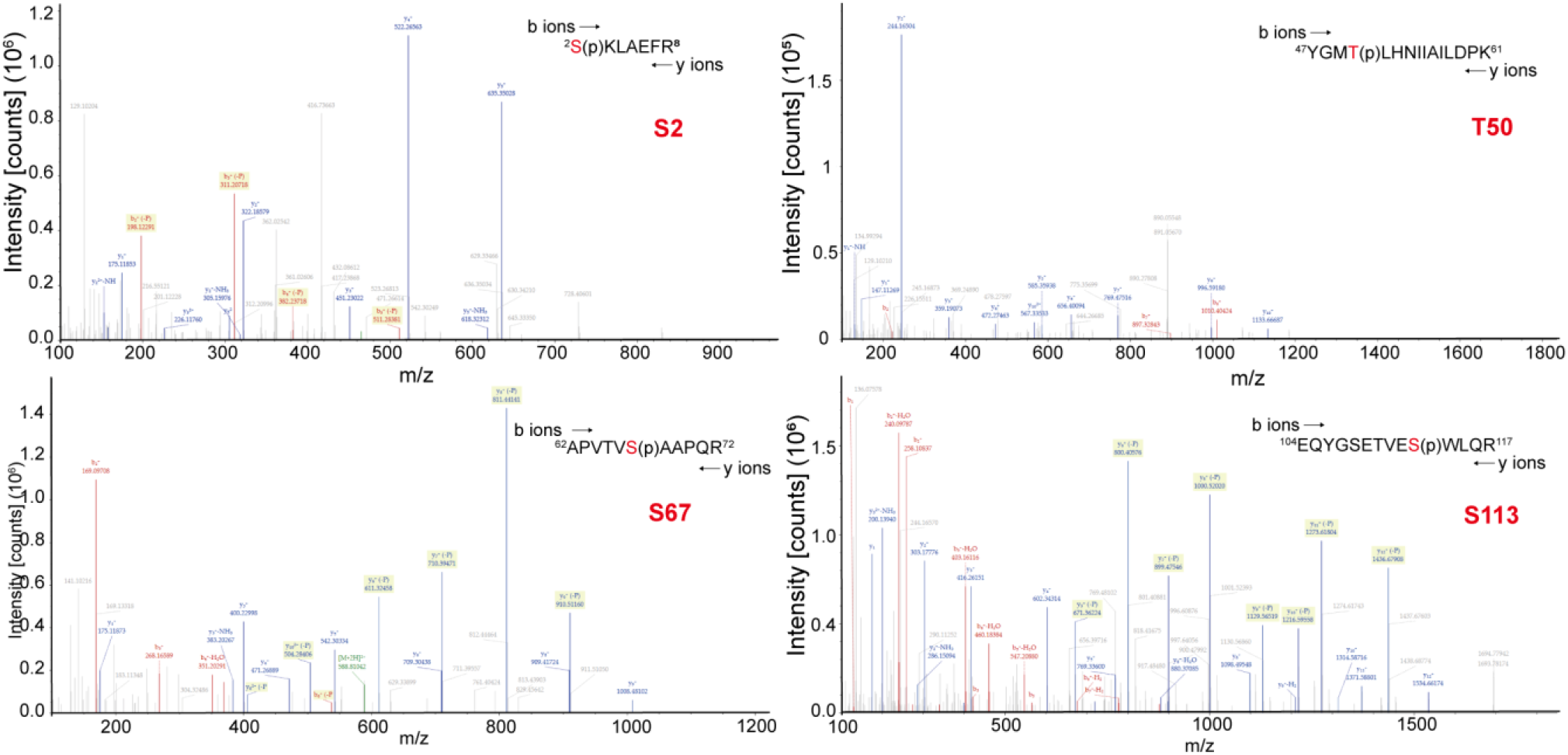
MvaU is phosphorylated at multiple sites when PfkA is expressed in PAO1. Phosphorylation sites of MvaU identified when PfkA is expressed in PAO1. MvaU-His protein expressed from pHERD20T-*mvaU-*His*-pfkA* in PAO1 was purified and subjected to LC–MS/MS analysis after trypsin digestion. Four phosphorylation sites in MvaU-His were identified, including S2, T50, S67 (S65 and S67 are too close to distinguish) and S113.

**Extended Data Fig. 5.**
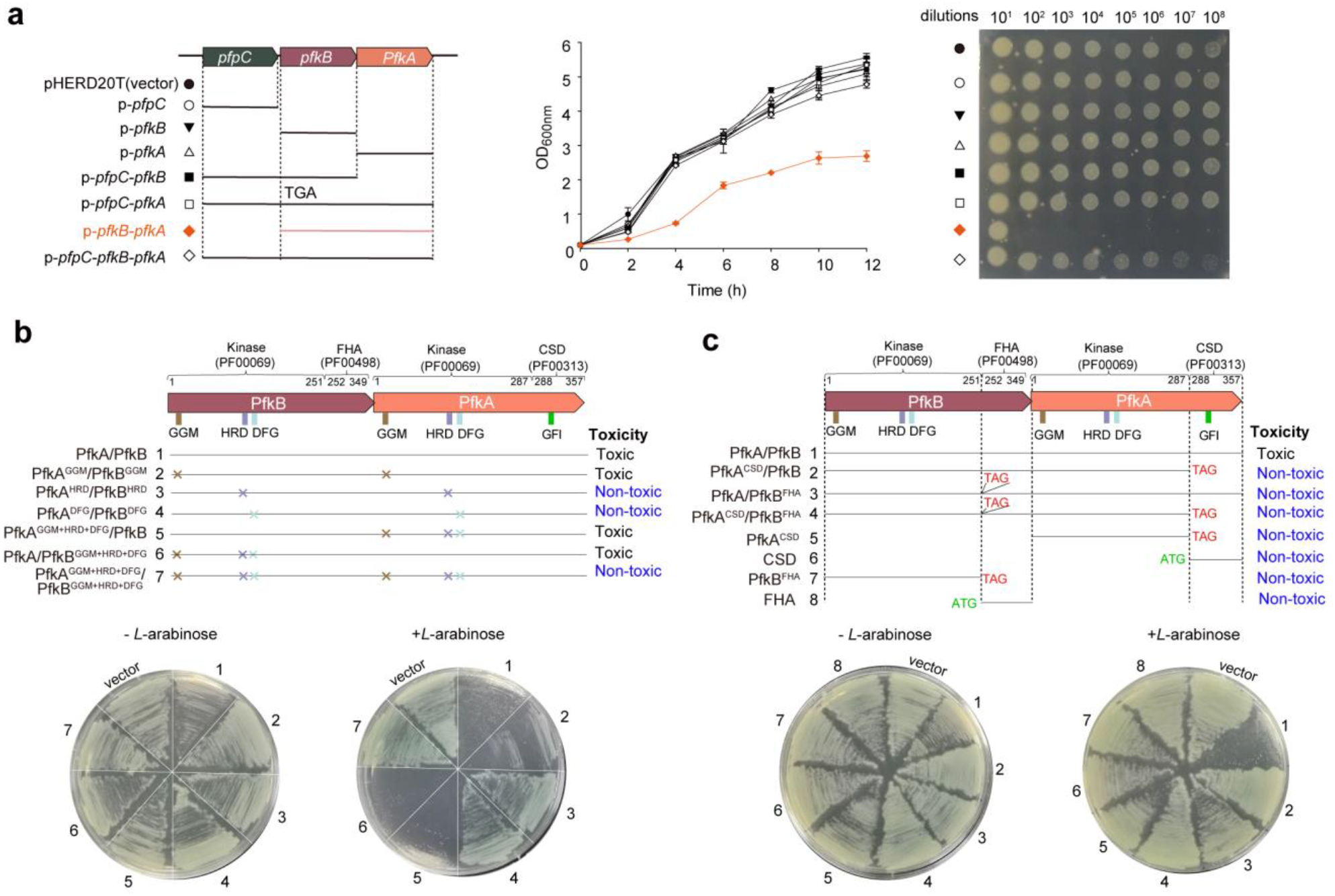
MPAO1 KKP_MP_ constitutes a tripartite TA system. **a**, Left: schematic of combinatorial introduction of KKP_MP_ components via pHERD20T into PAO1 *P. aeruginosa*. TGA indicates where the start codon of PfkB was replaced by TGA to prevent PfkB translation. Toxicity of introduced KKP_MP_ components was assessed with liquid growth (middle) media-based assay with *L*-arabinose for 12 h and CFU were assessed at 6 h (right). **b**, Toxicity of KKP_MP_ components with two or three conserved kinase motifs in PfkA and/or PfkB mutated at the N-terminus. Mutating two HRDs or two DFGs in both PfkA and PfkB abolished toxicity. **c**, Toxicity of KKP_MP_ components containing truncated PfkA and PfkB. The C-terminus of PfkA or PfkB is important for toxicity. TAG and ATG indicate insertion of a stop codon or start codon, respectively. Three independent biological replicates are tested and only representative images were shown for **a, b, c**.

**Extended Data Fig. 6.**
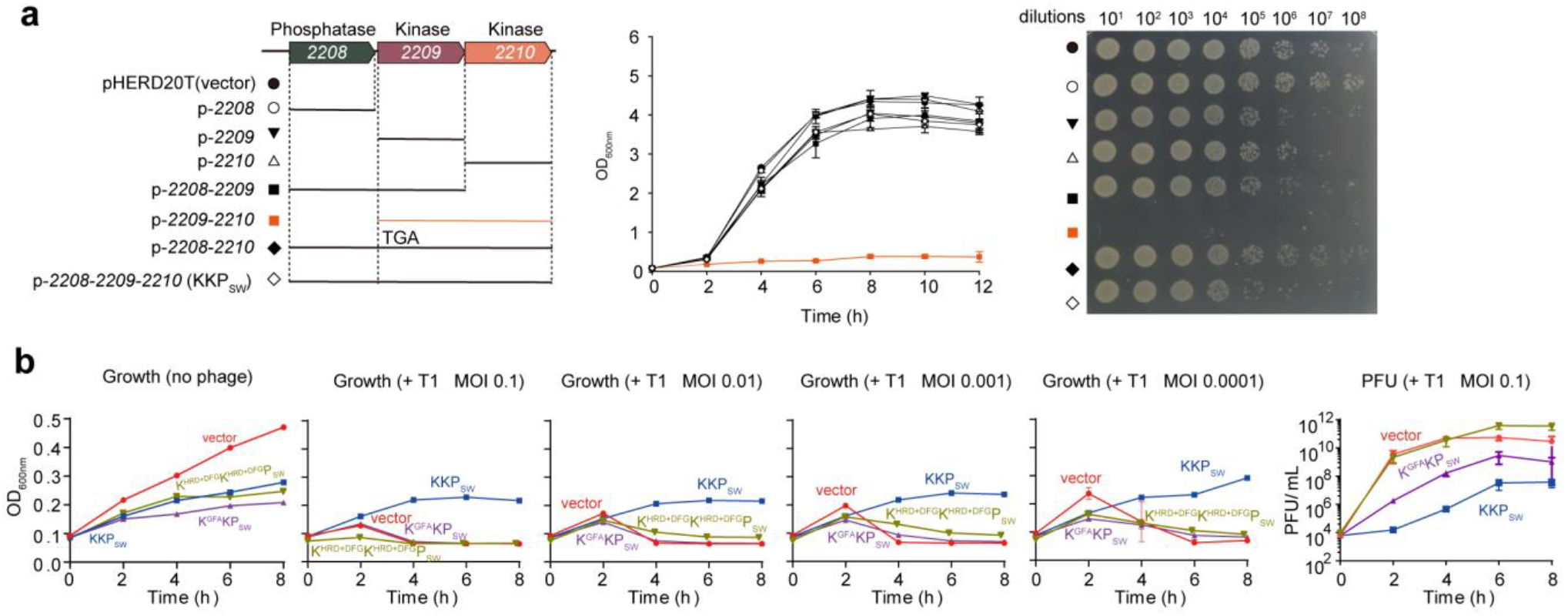
KKP_SW_ from the P4 prophage of *Shewanella* provides defense against lytic phages. **a**, Left: schematics of combinatorial introduction of KKP_SW_ components via pHERD20T into *E. coli* K12. Toxicity of introduced KKP_SW_ components was assessed with liquid growth (middle) media-based assays with *L*-arabinose for 12 h and CFU were assessed at 6 h (right). **b**, Growth curves for *E. coli* K12 harboring an empty vector or plasmid-encoded KKP_SW_, K^GFA^KP_SW_ and K^HRD+DFG^K^HRD+DFG^P_SW_ after infection with phage T1 at varying MOIs. Pfu was determined at indicated time points after infection with T1 at MOI of 0.1 (shown in the right panel). Three independent biological replicates are tested and data are shown as the mean ± SD.

**Extended Data Fig. 7.**
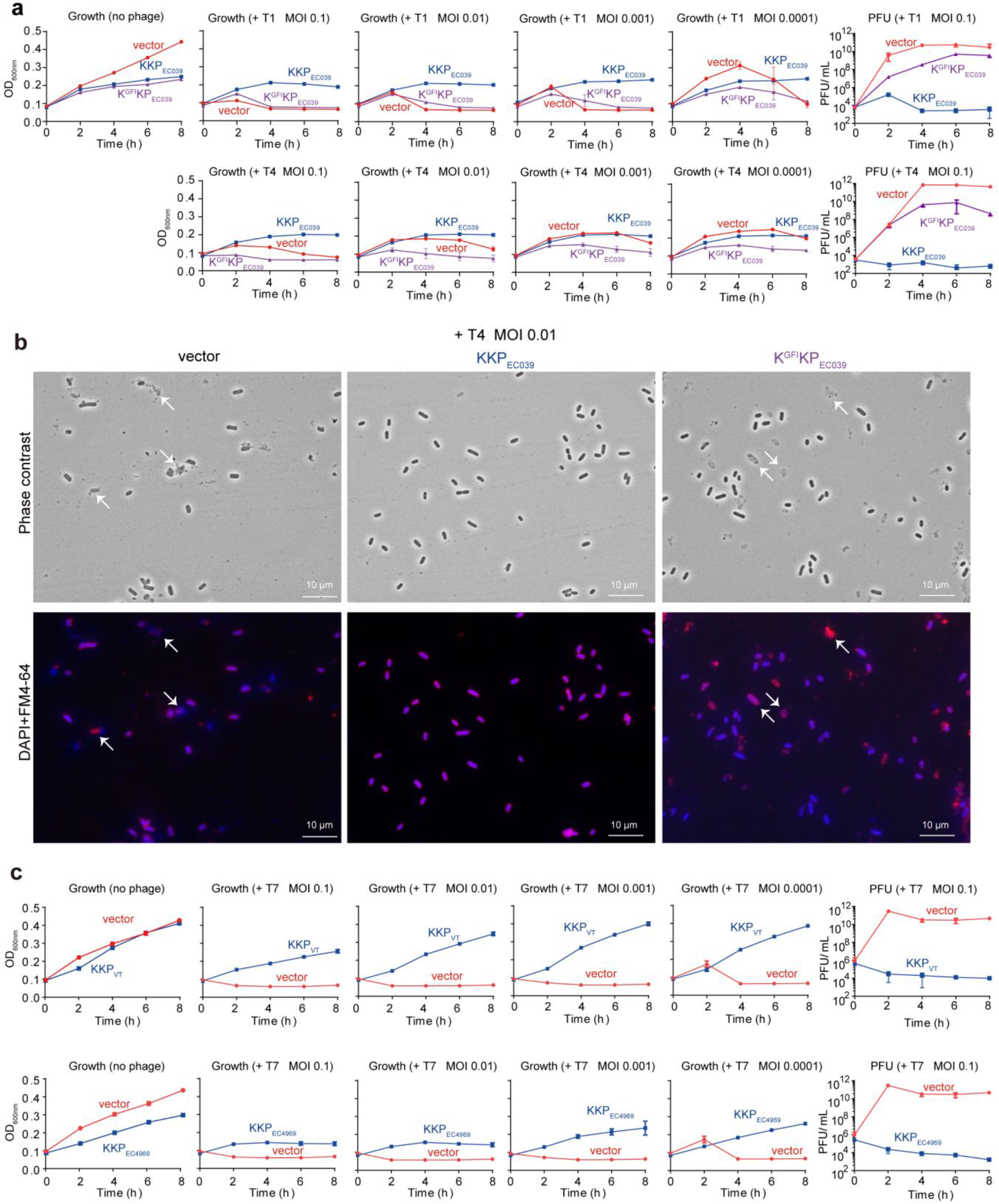
KKP modules from *E. coli* and *Vibrio* prophages provide defense against lytic phages. **a**, Growth curves for *E. coli* K12 harboring an empty vector or plasmid-encoded KKP_EC039_ or K^GFI^KP_EC039_ from the P2 prophage of *E. coli* 15EC039 after infection with T1 or T4 phages at varying MOI. Pfu was determined at the indicated time points after infection with T1 at MOI of 0.1 (shown in the right panel). **b**, Microscopy of FM4-64 and DAPI-stained cells after infection with T4 for 3 h at MOI of 0.01. Arrows indicated lysed cells. **c**, Growth curves for *E. coli* K12 harboring an empty vector or plasmid-encoded KKP_VT_ from P2 prophage of *V. tasmaniensis* or KKP_EC4969_ from P2 prophage of *E. coli* MPEC4969 after infection with T7 phages at varying MOIs. For these experiments, at least three independent biological replicates are used and data are shown as the mean ± SD.

