## Supplementary material for "Dual control of lysogeny and phage defense by a phosphorylation-based toxin/antitoxin system"

**This PDF file includes:**

Tables S1, S2, S4, S5, S8 and S9

References

**Legends for the Tables S3, S6 and S7 which are uploaded as separate excel files:**

**Table S3**. Published articles with sequencing raw data deposited in the NCBI SRA database. Sequencing reads were mapped to the reference genome of MPAO1 (CP079712). Average depths were calculated of Pf4 and Pf6 genome regions as well as the upstream and downstream 1000 bp regions flanking the prophage genomes. Y indicates yes and N indicates no.

**Table S6**. Distribution of KKP modules in bacteria. Data were retrieved from the IMG/G database. Gene IDs were indicated.

**Table S7.** List of phosphorylation sites in PAO1 genome that are modified by $KK_{MP}$ or $KKP_{MP}$ identified by phosphoproteome.

**Table S1**. **List of the Pf4 and Pf6 prophage genes in MPAO1 and their locations in PAO1 genome**. The core prophage genes are highlighted in bold letters.

| Pf4 in MPAO1 (12416 bp) | | | | | | Pf6 in MPAO1 (12148 bp) | | | | | Nucleotide | | Protein | |
|---|---|---|---|---|---|---|---|---|---|---|---|---|---|---|
| Locus tag (MPAO1) | Locus tag (PAO1) | Gene name | Start | Stop | Strand | Locus tag (MPAO1) | Gene name | Start | Stop | Strand | Coverage | Identity | Coverage | Identity |
| | | *attR* | 4717479 | 4717505 | - | | | | | | | | | |
| MP_4487 | PA0729 | *pfiT* | 4717628 | 4717975 | - | | | | | | | | | |
| MP_4488 | PA0728.1 | *pfiA* | 4717985 | 4718236 | - | | *attR* | 5241438 | 5241519 | | | | | |
| MP_4489 | PA0728 | *intF4* | 4718450 | 4719433 | - | MP_4975 | *intF6* | 5241680 | 5242681 | - | 0 | 0 | 98% | 40.66% |
| **MP_4490** | **PA0727** | ***repF4*** | **4719433** | **4720725** | **-** | **MP_4976** | ***repF6*** | **5242678** | **5243889** | **-** | **96%** | **92.30%** | **93%** | **95.29%** |
| **MP_4491** | **PA0726.1** | | **4720857** | **4720958** | **-** | **MP_4977** | | **5244102** | **5244203** | **-** | **100%** | **97.06%** | **100%** | **93.94%** |
| **MP_4492** | **PA0726** | | **4720955** | **4722229** | **-** | **MP_4978** | | **5244200** | **5245474** | **-** | **100%** | **98.27%** | **100%** | **97.88%** |
| **MP_4493** | **PA0725** | ***gVI*** | **4722233** | **4722589** | **-** | **MP_4979** | ***gVI*** | **5245478** | **5245834** | **-** | **100%** | **99.16%** | **100%** | **100%** |
| **MP_4494** | **PA0724** | ***gIII*** | **4722594** | **4723856** | **-** | **MP_4980** | ***gIII*** | **5245839** | **5247098** | **-** | **100%** | **98.10%** | **100%** | **98.81%** |
| **MP_4495** | **PA0723** | ***gVIII*** | **4723992** | **4724240** | **-** | **MP_4981** | ***gVIII*** | **5247234** | **5247482** | **-** | **100%** | **99.20%** | **100%** | **100%** |
| **MP_4496** | **PA0722** | ***gIX*** | **4724253** | **4724504** | **-** | **MP_4982** | ***gIX*** | **5247495** | **5247746** | **-** | **100%** | **100%** | **100%** | **100%** |
| **MP_4497** | **PA0721** | ***gVII*** | **4724517** | **4724609** | **-** | **MP_4983** | ***gVII*** | **5247759** | **5247851** | **-** | **100%** | **97.85%** | **100%** | **96.67%** |
| **MP_4498** | **PA0720** | | **4724626** | **4725060** | **-** | **MP_4984** | | **5247868** | **5248302** | **-** | **100%** | **96.09%** | **100%** | **97.92%** |
| **MP_4499** | **PA0719** | | **4725195** | **4725572** | **-** | **MP_4985** | | **5248469** | **5248813** | **-** | **99%** | **97.38%** | **79%** | **98.99%** |
| **MP_4500** | **PA0718** | | **4725576** | **4725866** | **-** | **MP_4986** | | **5248817** | **5249107** | **-** | **100%** | **96.91%** | **100%** | **96.88%** |
| **MP_4501** | **PA0717** | | **4725870** | **4726082** | **-** | **MP_4987** | | **5249111** | **5249239** | **-** | **60%** | **97.67%** | **60%** | **97.62%** |
| MP_4502 | PA0716.2 | *xisF4* | 4726084 | 4726299 | - | MP_4988 | *xisF6* | 5249319 | 5249549 | - | 0 | 0 | 49% | 34.29% |
| MP_4503 | PA0716.1 | *pf4r* | 4726418 | 4726684 | + | MP_4989 | *pf6r* | 5249717 | 5249986 | + | 0 | 0 | 100% | 50.56% |
| MP_4504 | PA0716 | | 4726973 | 4728298 | - | MP_4990 | *pfpC* | 5250121 | 5251218 | + | | | | |
| MP_4505 | PA0715 | | 4728301 | 4729257 | - | MP_4991 | *pfkB* | 5251221 | 5252270 | + | | | | |
| MP_4506 | PA0714.1 | *phrD* | 4729635 | 4729707 | - | MP_4992 | *pfkA* | 5252267 | 5253340 | + | | | | |
| | | *attL* | 4729868 | 4729894 | - | | *attL* | 5253510 | 5253589 | | | | | |

**Table S2.** The presence of Pf4 and Pf6 loci in the selected *P. aeruginosa* strains were analyzed by BLASTN. The four genomes used in Fig. 2a are shown in blue fonts and two ATCC strains were shown in bold fonts. In addition, the strains with determined sources summarized by C. E., Chandler et al.[1] were also analyzed, and the assembly of these strains were downloaded from NCBI under BioProject accession number PRJNA490649. Y indicates yes and N indicates no.

| GenBank/Assembly ID | Strain name | Pf4 | Pf6 | Source | Date acquired | PubMed ID | Date of publish |
|---|---|---|---|---|---|---|---|
| **MPAO1 sublines** | | | | | | | |
| **CP079712** | **MPAO1** | **Y** | **Y** | **McDermott, University of Washington** | **07/2020** | **this study** | |
| **CP027857** | **MPAO1 (C. Manoil)** | **Y** | **Y** | **C. Manoil, University of Washington** | **2017** | **33127897** | 2020 Oct 30 |
| GCA_003957075 | MPAO-1 (C. Manoil) | Y | Y | C. Manoil, University of Washington | 01/2003 | 30530517 | 2019 Feb 11 |
| GCA_003957195 | MPAO-1 (C. Manoil) | Y | Y | C. Manoil, University of Washington | 08/2008 | 30530517 | 2019 Feb 11 |
| GCA_003957175 | PAO-1 (B. Iglewski) | Y | Y | E.P. Greenberg, University of Washington | 10/2000 | 30530517 | 2019 Feb 11 |
| CP050052 | PAO1 (B. Iglewski) | Y | Y | E. P. Greenberg, University of Washington | 04/2019 | unpublished | |
| GCF_003957105 | H103 PAO-1 AK957 | Y | Y | R. Hancock, University of British Columbia | 02/2000 | 30530517 | 2019 Feb 11 |
| GCA_003957205 | PAO-1 (B. Holloway) | Y | Y | D. Ohman, Virginia Commonwealth University | 05/2001 | 30530517 | 2019 Feb 11 |
| GCA_003957095 | PAO-1 V | Y | Y | J. Goldberg, University of Virginia | 07/2003 | 30530517 | 2019 Feb 11 |
| GCA_003957085 | PAO-1 | Y | Y | H. Nikaido, University of California, Berkeley | 08/2003 | 30530517 | 2019 Feb 11 |
| GCA_003957115 | PAO-1 | Y | Y | A. Prince, Columbia University | 11/2003 | 30530517 | 2019 Feb 11 |
| CP050054/CP050053 | PAO1 | Y | Y | Y. Liu, South China Agricultural University | 04/2018 | unpublished | |
| CP053028 | PAO1 | Y | Y | S.P. Diggle, Georgia Institute of Technology | 01/2014 | 32341475 | 2020 Mar 23 |
| not available | PAO1 | Y | Y | Scott A. Rice, Nanyang Technological University | not available | 34452479 | 2021 Aug 15 |
| LN871187 | PAO1_Orsay1 | Y | Y | Pourcel, Institut de Genetique et Microbiologie | not available | unpublished | |
| CP032126 | PAO1161 | Y | Y | A. Kawalek, Institute for Biochemistry and Biophysics PAS | 11/2017 | 31906858 | 2020 Jan 6 |
| **NZ_CP017149** | **PAO-1C (ATCC 15692)** | **Y** | **Y** | **ATCC (deposited by BW Holloway)** | **not available** | unpublished | |
| **CP041008** | **PAO1 (ATCC BAA-47)** | **Y** | **Y** | **ATCC (deposited by HW Ackermann)** | **not available** | unpublished | |
| **PAO1 sublines** | | | | | | | |
| **CP085082** | **PAO1** | **Y** | **N** | **J.J. Mekalanos, Harvard Medical School** | **10/2020** | **this study** | |
| **AE004091** | **PAO1** | **Y** | **N** | **P. Phibbs, University of Georgia** | **not available** | **10984043** | **2000 Aug 31** |
| GCA_003957185 | PAO-1 | Y | N | J. Burns, University of Washington | 02/2000 | 30530517 | 2019 Feb 11 |
| CP006832/CP006831 | PAO1 | Y | N | H.W.D., Yu, Marshall University | not available | 24336371 | 2013 Dec 12 |
| CP006705 | PAO1 (PA0581) | Y | N | H.W.D., Yu, Marshall University | not available | 24115549 | 2013 Oct 10 |
| CP053110 to CP053119 | PAO1 | Y | N | S. Fong, University of Adelaide | 12/2017 | unpublished | |
| CP047061 to CP047068 | PAO1 | Y | N | Balint Csorgo, UCSF | not available | unpublished | |
| CP032540/CP032541 | PAO1 | Y | N | Progenesis Technologies, LLC | 09/2014 | unpublished | |
| CP034908 | PA0750 | Y | N | H.P. Schweizer, Colorado State University | 2005 | 17005832 | 2006 Oct |

**Table S4**. **RNA-seq reads of each Pf4 and Pf6 genes in the biofilm cells and in the planktonic cells of MPAO1 strain.** Three replicates were used and shown. NA indicates not available. FC, CPM and FDR indicate fold change, counts per million and false discovery rate, respectively.

| **Locus tag (MPAO1)** | **Gene product** | **Planktonic cells** | | | **Biofilm cells** | | | **LogFC** | **LogCPM** | **FDR** |
|---|---|---|---|---|---|---|---|---|---|---|
| | | **# 1** | **# 2** | **# 3** | **# 1** | **# 2** | **# 3** | | | |
| **Pf4 prophage** | | | | | | | | | | |
| **MP_4487** | PfiT | 59.0 | 69.9 | 45.6 | 583.1 | 272.6 | 468.9 | 3.0 | 6.4 | 5.29E-36 |
| **MP_4488** | PfiA | 60.2 | 47.0 | 63.6 | 539.2 | 302.2 | 369.8 | 2.9 | 5.9 | 1.55E-24 |
| **MP_4489** | IntF4 | 1.7 | 4.0 | 2.3 | 459.7 | 103.6 | 155.5 | 6.5 | 6.8 | 2.72E-44 |
| **MP_4490** | RepF4 | 2.9 | 3.8 | 3.6 | 892.8 | 196.0 | 316.5 | 7.1 | 8.2 | 6.83E-54 |
| **MP_4491** | Hypothetical | 0.0 | 0.0 | 0.0 | 41.5 | 1.8 | 9.3 | 6.3 | 0.1 | 0.00088 |
| **MP_4492** | Hypothetical | 1.0 | 0.6 | 1.0 | 576.3 | 122.4 | 177.4 | 8.3 | 7.5 | 8.38E-53 |
| **MP_4493** | pVI | 0.0 | 0.9 | 0.0 | 23.7 | 3.0 | 0.5 | 4.7 | 0.8 | 0.001108 |
| **MP_4494** | pIII | 0.3 | 1.2 | 1.3 | 1196.3 | 249.6 | 426.0 | 9.4 | 8.5 | 4.14E-62 |
| **MP_4495** | pVIII | 1.3 | 4.3 | 6.4 | 11773.3 | 1518.9 | 1889.6 | 10.2 | 9.2 | 2.20E-35 |
| **MP_4496** | pIX | 0.0 | 0.0 | 0.0 | 0.0 | 0.0 | 0.0 | NA | NA | NA |
| **MP_4497** | pVII | 0.0 | 0.0 | 0.0 | 123.4 | 15.5 | 34.5 | 7.9 | 1.4 | 1.12E-08 |
| **MP_4498** | Hypothetical | 8.3 | 23.8 | 9.2 | 20618.8 | 4612.1 | 8445.6 | 9.7 | 11.2 | 5.79E-77 |
| **MP_4499** | Hypothetical | 8.6 | 9.7 | 0.9 | 9635.9 | 2351.2 | 4761.3 | 9.8 | 10.0 | 1.07E-71 |
| **MP_4500** | Hypothetical | 3.4 | 4.7 | 8.8 | 5423.0 | 2356.9 | 3570.8 | 9.5 | 9.1 | 8.37-126 |
| **MP_4501** | Hypothetical | 0.8 | 0.7 | 3.0 | 1681.4 | 588.2 | 744.9 | 9.4 | 6.7 | 7.87E-83 |
| **MP_4502** | XisF4 | 2.3 | 1.4 | 2.2 | 5079.5 | 950.1 | 1656.8 | 10.3 | 8.0 | 1.58E-62 |
| **MP_4503** | Pf4r | 24.4 | 23.9 | 18.6 | 48.3 | 58.0 | 110.9 | 1.9 | 3.7 | 1.36E-06 |
| **MP_4504** | Hypothetical | 48.2 | 38.2 | 32.7 | 39.9 | 68.0 | 94.3 | 0.99 | 6.2 | 0.01418 |
| **MP_4505** | Hypothetical | 48.4 | 22.9 | 23.4 | 33.9 | 60.2 | 91.3 | 1.2 | 5.6 | 0.01255 |
| **MP_4506** | PhrD | 464.6 | 878.3 | 875.3 | 391.6 | 254.0 | 434.2 | -0.92 | 5.3 | 0.00379 |
| **Pf6 prophage** | | | | | | | | | | |
| **MP_4975** | IntF6 | 4.9 | 2.3 | 3.8 | 61.9 | 93.2 | 142.0 | 4.9 | 5.8 | 9.62E-27 |
| **MP_4976** | RepF6 | 3.2 | 2.5 | 2.1 | 164.9 | 237.5 | 393.0 | 6.9 | 7.5 | 4.69E-49 |
| **MP_4977** | Hypothetical | 0.0 | 0.0 | 0.0 | 63.2 | 88.2 | 160.9 | 9.1 | 2.5 | 1.01E-18 |
| **MP_4978** | Hypothetical | 0.1 | 0.6 | 0.5 | 70.1 | 135.1 | 200.5 | 8.6 | 6.6 | 1.07E-44 |
| **MP_4979** | pVI | 0.0 | 0.0 | 0.0 | 10.7 | 9.1 | 9.5 | 7.5 | 1.0 | 2.15E-09 |

| | | | | | | | | | | |
|---|---|---|---|---|---|---|---|---|---|---|
| **MP_4980** | pIII | 1.0 | 0.5 | 1.0 | 189.8 | 335.9 | 525.1 | 8.9 | 7.9 | 1.92E-52 |
| **MP_4981** | pVIII | 0.0 | 1.2 | 2.6 | 3837.9 | 2455.9 | 2303.2 | 11.2 | 8.5 | 8.44E-126 |
| **MP_4982** | pIX | 0.0 | 0.0 | 0.0 | 0.0 | 0.0 | 0.0 | NA | NA | NA |
| **MP_4983** | pVII | 0.0 | 0.0 | 0.0 | 10.8 | 7.7 | 24.3 | 6.0 | -0.1 | 0.000402 |
| **MP_4984** | Hypothetical | 4.1 | 6.3 | 4.4 | 4002.7 | 4457.3 | 6895.2 | 10.2 | 10.2 | 1.91E-110 |
| **MP_4985** | Hypothetical | 3.8 | 4.4 | 0.9 | 4505.9 | 4766.5 | 8160.2 | 11.1 | 10.1 | 4.38E-108 |
| **MP_4986** | Hypothetical | 0.6 | 1.0 | 2.2 | 2020.1 | 2039.6 | 3080.0 | 11.0 | 8.5 | 8.22E-108 |
| **MP_4987** | Hypothetical | 2.5 | 0.0 | 2.5 | 1373.3 | 1618.7 | 1675.9 | 10.0 | 6.8 | 4.59E-81 |
| **MP_4988** | XisF6 | 3.5 | 4.6 | 1.4 | 4576.2 | 6590.4 | 10843.9 | 11.4 | 9.9 | 1.21E-85 |
| **MP_4989** | Pf6r | 186.0 | 146.3 | 99.1 | 1427.1 | 462.0 | 666.6 | 2.6 | 7.0 | 8.03E-17 |
| **MP_4990** | PfpC | 93.1 | 67.4 | 46.4 | 468.6 | 209.2 | 306.5 | 2.3 | 7.8 | 1.02E-16 |
| **MP_4991** | PfkB | 162.3 | 109.5 | 76.3 | 2198.4 | 953.6 | 1415.6 | 3.8 | 9.7 | 3.64E-34 |
| **MP_4992** | PfkA | 731.0 | 674.1 | 431.5 | 933.8 | 319.8 | 466.5 | -0.02 | 9.3 | 0.97 |

**Table S5. List of proteins that are modified by PfkA kinase identified by phosphoproteome.** The fold changes indicated the phosphorylation levels of proteins controlled by PfkA via pHERD20T-*pfkA* overproduction versus those in cells harboring pHERD20T vector. Two replicates were used, and those proteins with fold changes > 2.5 and coefficient of variation (CV) < 0.1) were selected.

| Locus tag (MPAO1) | Locus tag (PAO1) | Protein | Position | Fold change | CV value | Description |
|---|---|---|---|---|---|---|
| MP_2627 | PA2480 | | 391T | 4.554 | 0.006 | Signal transduction histidine kinase |
| MP_4036 | PA1155 | NrdB | 408T | 3.968 | 0.008 | Ribonucleotide-diphosphate reductase subunit beta |
| MP_618 | PA0577 | DnaG | 527T | 3.816 | 0.088 | DNA primase |
| MP_1549 | PA3481 | ApbC | 353S | 3.777 | 0.022 | ATPases involved in chromosome partitioning |
| MP_3583 | PA1585 | SucA | 62T | 3.616 | 0.072 | Dehydrogenase (E1) component related enzymes |
| MP_82 | PA0074 | PpkA | 270T | 3.261 | 0.087 | Ser/Thr kinase |
| MP_723 | PA4274 | RplK | 71T | 3.101 | 0.046 | Ribosomal protein L11 |
| MP_5588 | PA5239 | Rho | 84S | 3.010 | 0.060 | Transcription termination factor |
| MP_2467 | PA2629 | PurB | 296S | 2.975 | 0.088 | Adenylosuccinate lyase |
| **MP_2427** | **PA2667** | **MvaU** | **65T/67S** | **2.892** | **0.019** | **H-NS family protein** |
| MP_5084 | PA4759 | DapB | 207T | 2.640 | 0.069 | Dihydrodipicolinate reductase |
| MP_4426 | PA0787 | | 79S | 2.624 | 0.089 | Predicted ATPase |
| MP_5631 | PA5279 | | 49S | 2.609 | 0.085 | Uncharacterized protein conserved in bacteria |
| MP_5697 | PA5343 | | 246S | 2.604 | 0.026 | Nucleoside-diphosphate-sugar epimerases |
| MP_5797 | PA5435 | PycB | 462T | 2.517 | 0.022 | Pyruvate/oxaloacetate carboxyltransferase |
| MP_1500 | PA3529 | | 198S | 0.229 | 0.049 | Peroxidase |
| MP_1153 | PA3861 | RhlB | 38S | 0.129 | 0.044 | ATP-dependent RNA helicase RhlB |
| MP_180 | PA0170 | | 68T | 0.017 | 0.083 | DUF1987 domain-containing protein |

**Table S8. Bacterial strains and plasmids used in this study.** R indicates resistance, Gm indicates gentamycin, Amp indicates ampicillin, MP indicates MPAO1 strain. KKP indicates the *pfkA-pfkB-pfpC* cascade.

| | Description | Source |
|---|---|---|
| **Strains** | | |
| PAO1 | wild-type | [2] |
| MPAO1 | wild-type | [3] |
| MP-ΔPf4 | whole Pf4 prophage removed from MPAO1 host chromosome | [3] |
| MP-ΔPf6core | the core genes from MP-4976 to MP_4987 removed from MPAO1 host chromosome | this study |
| MP-ΔPf4ΔPf6core | whole Pf4 prophage removed from MP-ΔPf6core host chromosome | this study |
| pVIII::GFP | *gVIII* gene encoding Pf4 major coat protein pVIII in-frame fused with *gfp* gene in PAO1 host chromosome | this study |
| MP-pVIII::GFP | *gVIII* gene encoding Pf4 major coat protein pVIII in-frame fused with *gfp* gene in MPAO1 host chromosome | this study |
| MP-Δ*mvaU*Δ*mvaT* | *mvaT* and *mvaU* double deletion mutant derived from MPAO1 | [3] |
| PAO1::*pilC*$_{MP}$ | *pilC* gene from MPAO1 was integrated into Δ*pilC* host chromosome | this study |
| PAO1::*pilC*$_{MP}$::KKP$_{MP}$ | the KKP$_{MP}$ cascades were integrated into PAO1::*pilC*$_{MP}$ host chromosome | this study |
| PAO1::KKP$_{MP}$ | the KKP$_{MP}$ cascades were integrated into PAO1 host chromosome | this study |
| PAO1::*pilC*$_{MP}$::K$^{GFI}$KP$_{MP}$ | the mutant KKP$_{MP}$ cascade (the 307$^{th}$ -309$^{th}$ aa codon GGATTCATT in *pfkA* was mutated to GACTATGCA) were integrated into PAO1::*pilC*$_{MP}$ host chromosome | this study |
| WM3064 | *thrB*1004 *pro thi rpsL hsdS lacZ*ΔM15 RP4-1360) Δ(*araBAD*)567 Δ*dapA*1341::[*erm pir(*wt)] | W. Metcalf, UIUC |
| MG655 | K-12 *F*$^{-}$ *lambda*$^{-}$ *ilvG- rfb-50 rph-1* | lab stored |
| **Plasmids** | | |
| pEX18Ap | Ap$^{R}$, *oriT*$^{+}$, *sacB*$^{+}$, gene replacement vector | [4] |
| pEX18Gm | Gm$^{R}$, *oriT*$^{+}$, *sacB*$^{+}$, gene replacement vector | [4] |
| pFLP2 | Ap$^{R}$, Flp recombinase-expressing plasmid | [4] |
| pPS856 | Ap$^{R}$, Gm$^{R}$; for amplifying gentamycin resistance cassette | [4] |
| pEX18Gm-ΔPf6core-up-down | Gm$^{R}$, Ap$^{R}$, for deleting Pf6core region | this study |
| pEX18Gm-up-pVIII::GFP-down | Gm$^{R}$, Ap$^{R}$, for Pf4 *gVIII* in-frame fused with *gfp* gene in MPAO1 host chromosome | this study |
| pEX18Gm-*pilC*-up-down | Gm$^{R}$, Ap$^{R}$, for deleting *pilC* in PAO1 host chromosome | this study |
| pEX18Gm-up-*pilC*$_{MP}$-down | Gm$^{R}$, Ap$^{R}$, for integration of *pilC*$_{MP}$ gene from MPAO1 into Δ*pilC* host chromosome | this study |
| pEX18Gm-up-Gm-KKP-down | Gm$^{R}$, Ap$^{R}$, for integration of KKP$_{MP}$ cascade from MPAO1 into PAO1::*pilC*$_{MP}$ host chromosome | this study |
| pEX18Gm-up-Gm-K$^{GFI}$KP-down | Gm$^{R}$, Ap$^{R}$, for integration of K$^{GFI}$KP$_{MP}$ cascade mutant into PAO1:: *pilC*$_{MP}$ host chromosome | this study |
| pHERD20T | Ap$^{R}$, expression vector with araC-P$_{BAD}$ promoter | [5] |
| pHERD20T-*mvaU* | Ap$^{R}$, *mvaU* in pHERD20T NcoI/HindIII | this study |
| pHERD20T-*mvaU*$^{S2A}$ | Ap$^{R}$, the second aa codon TCC in *mvaU* was mutated to GCC in pHERD20T-*mvaU* | this study |

| | | |
|---|---|---|
| pHERD20T-*mvaU*$^{S2D}$ | Ap$^{R}$, the second aa codon TCC in *mvaU* was mutated to GAC in pHERD20T-*mvaU* | this study |
| pHERD20T-*mvaU*$^{S26A}$ | Ap$^{R}$, the 26$^{th}$ aa codon AGC in *mvaU* was mutated to GCC in pHERD20T-*mvaU* | this study |
| pHERD20T-*mvaU*$^{S26D}$ | Ap$^{R}$, the 26$^{th}$ aa codon AGC in *mvaU* was mutated to GAC in pHERD20T-*mvaU* | this study |
| pHERD20T-*mvaU*$^{T50A}$ | Ap$^{R}$, the 50$^{th}$ aa codon ACC in *mvaU* was mutated to GCC in pHERD20T-*mvaU* | this study |
| pHERD20T-*mvaU*$^{T50D}$ | Ap$^{R}$, the 50$^{th}$ aa codon ACC in *mvaU* was mutated to GAC in pHERD20T-*mvaU* | this study |
| pHERD20T-*mvaU*$^{T65A}$ | Ap$^{R}$, the 65$^{th}$ aa codon ACC in *mvaU* was mutated to GCC in pHERD20T-*mvaU* | this study |
| pHERD20T-*mvaU*$^{T65D}$ | Ap$^{R}$, the 65$^{th}$ aa codon ACC in *mvaU* was mutated to GAC in pHERD20T-*mvaU* | this study |
| pHERD20T-*mvaU*$^{S67A}$ | Ap$^{R}$, the 67$^{th}$ aa codon AGC in *mvaU* was mutated to GCC in pHERD20T-*mvaU* | this study |
| pHERD20T-*mvaU*$^{S67D}$ | Ap$^{R}$, the 67$^{th}$ aa codon AGC in *mvaU* was mutated to GAC in pHERD20T-*mvaU* | this study |
| pHERD20T-*mvaU*$^{S108A}$ | Ap$^{R}$, the 108$^{th}$ aa codon TCC in *mvaU* was mutated to GCC in pHERD20T-*mvaU* | this study |
| pHERD20T-*mvaU*$^{S108D}$ | Ap$^{R}$, the 108$^{th}$ aa codon TCC in *mvaU* was mutated to GAC in pHERD20T-*mvaU* | this study |
| pHERD20T-*mvaU*$^{S113A}$ | Ap$^{R}$, the 113$^{th}$ aa codon TCC in *mvaU* was mutated to GCC in pHERD20T-*mvaU* | this study |
| pHERD20T-*mvaU*$^{S113D}$ | Ap$^{R}$, the 113$^{th}$ aa codon TCC in *mvaU* was mutated to GAC in pHERD20T-*mvaU* | this study |
| pHERD20T-*mvaU*-His-*pfkA* | Ap$^{R}$, N terminal His tagged *mvaU* was coexpressed with *pfkA* in pHERD20T NcoI/HindIII | this study |
| pHERD20T-*mvaU*-His-*pfkB* | Ap$^{R}$, N terminal His tagged *mvaU* was coexpressed with *pfkB* in pHERD20T NcoI/HindIII | this study |
| pHERD20T-*mvaU*-His-*pfkA-pfkB* | Ap$^{R}$, N terminal His tagged *mvaU* was coexpressed with *pfkA-pfkB* in pHERD20T NcoI/HindIII | this study |
| pHERD20T-*mvaU*-His-KKP$_{MP}$ | Ap$^{R}$, N terminal His tagged *mvaU* was coexpressed with KKP$_{MP}$ in pHERD20T NcoI/HindIII | this study |
| pHERD20T-*pfkA* | Ap$^{R}$, *pfkA* in pHERD20T NcoI/HindIII | this study |
| pHERD20T-*pfkA*$^{CSD}$ | Ap$^{R}$, mutant *pfkA* lacking CSD domain in pHERD20T NcoI/HindIII | this study |
| pHERD20T-*CSD* | Ap$^{R}$, CSD domain of *pfkA* in pHERD20T NcoI/HindIII | this study |
| pHERD20T-*pfkB* | Ap$^{R}$, *pfkB* in pHERD20T NcoI/HindIII | this study |
| pHERD20T-*pfkB*$^{FHA}$ | Ap$^{R}$, mutant *pfkB* lacking FHA domain in pHERD20T NcoI/HindIII | this study |
| pHERD20T-*FHA* | Ap$^{R}$, FHA domain of *pfkB* in pHERD20T NcoI/HindIII | this study |
| pHERD20T-*pfpC* | Ap$^{R}$, *pfpc* in pHERD20T NcoI/HindIII | this study |
| pHERD20T-*pfkA-pfkB* | Ap$^{R}$, *pfkA-pfkB* in pHERD20T NcoI/HindIII | this study |
| pHERD20T-*pfkA*$^{CSD}$-*pfkB* | Ap$^{R}$, mutant *pfkA-pfkB* without CSD domain in pHERD20T NcoI/HindIII | this study |
| pHERD20T-*pfkA-pfkB*$^{FHA}$ | Ap$^{R}$, mutant *pfkA-pfkB* without FHA domain in pHERD20T NcoI/HindIII | this study |
| pHERD20T-*pfkA*$^{CSD}$-*pfkB*$^{FHA}$ | Ap$^{R}$, mutant *pfkA-pfkB* without both CSD and FHA domains in pHERD20T NcoI/HindIII | this study |
| pHERD20T-*pfkA*$^{GGM}$-*pfkB* | Ap$^{R}$, mutant GGM motif in PfkA in *pfkA-pfkB* in pHERD20T NcoI/HindIII | this study |
| pHERD20T-*pfkA*$^{HRD}$-*pfkB* | Ap$^{R}$, mutant HRD motif in PfkA in *pfkA-pfkB* in pHERD20T NcoI/HindIII | this study |
| pHERD20T-*pfkA*$^{DFG}$-*pfkB* | Ap$^{R}$, mutant DFG motif in PfkA in *pfkA-pfkB* in pHERD20T NcoI/HindIII | this study |
| pHERD20T-*pfkA*$^{GFI}$-*pfkB* | Ap$^{R}$, mutant GFI motif in PfkA in *pfkA-pfkB* in pHERD20T NcoI/HindIII | this study |
| pHERD20T-*pfkA-pfkB*$^{GGM}$ | Ap$^{R}$, mutant GGM motif in PfkB in *pfkA-pfkB* in pHERD20T NcoI/HindIII | this study |

| | | |
|---|---|---|
| pHERD20T-*pfkA-pfkB*$^{HRD}$ | Ap$^R$, mutant HRD motif in PfkB in *pfkA-pfkB* in pHERD20T NcoI/HindIII | this study |
| pHERD20T-*pfkA-pfkB*$^{DFG}$ | Ap$^R$, mutant DFG motif in PfkB in *pfkA-pfkB* in pHERD20T NcoI/HindIII | this study |
| pHERD20T-*pfkA*$^{GGM}$-*pfkB*$^{GGM}$ | Ap$^R$, mutant GGM motifs in both PfkA and PfkB in *pfkA-pfkB* in pHERD20T NcoI/HindIII | this study |
| pHERD20T-*pfkA*$^{HRD}$-*pfkB*$^{HRD}$ | Ap$^R$, mutant HRD motifs in both PfkA and PfkB in *pfkA-pfkB* in pHERD20T NcoI/HindIII | this study |
| pHERD20T-*pfkA*$^{DFG}$-*pfkB*$^{DFG}$ | Ap$^R$, mutant DFG motifs in both PfkA and PfkB in *pfkA-pfkB* in pHERD20T NcoI/HindIII | this study |
| pHERD20T-*pfkA*$^{GGM+HRD+DFG}$-*pfkB* | Ap$^R$, the GGM, HRD and DFG motif in PfkA were mutated as PAO1::MP-*pilC*::K$^{GGM+HRD+DFG}$K$^{GGM+HRD+DFG}$P in *pfkA-pfkB,* and the generated *pfkA*$^{GGM+HRD+DFG}$-*pfkB* was ligated to pHERD20T NcoI/HindIII | this study |
| pHERD20T-*pfkA-pfkB*$^{GGM+HRD+DFG}$ | Ap$^R$, the GGM, HRD and DFG motif in PfkB were mutated as PAO1::MP-*pilC*::K$^{GGM+HRD+DFG}$K$^{GGM+HRD+DFG}$P in *pfkA-pfkB,* and the generated *pfkA*$^{GGM+HRD+DFG}$-*pfkB* was ligated to pHERD20T NcoI/HindIII | this study |
| pHERD20T-*pfkA*$^{GGM+HRD+DFG}$ -*pfkB*$^{GGM+HRD+DFG}$ | Ap$^R$, the GGM, HRD and DFG motif in both PfkA and PfkB were mutated as PAO1::MP-*pilC*::K$^{GGM+HRD+DFG}$K$^{GGM+HRD+DFG}$P in *pfkA-pfkB,* and the generated *pfkA*$^{GGM+HRD+DFG}$-*pfkB*$^{GGM+HRD+DFG}$ was ligated to pHERD20T NcoI/HindIII | this study |
| pHERD20T-*pfkA-pfpC* | Ap$^R$, *pfkA-pfpC* in pHERD20T NcoI/HindIII | this study |
| pHERD20T-*pfkB-pfpC* | Ap$^R$, *pfkB-pfpC* in pHERD20T NcoI/HindIII | this study |
| pHERD20T-*pfkA-pfkB-pfpC* | Ap$^R$, KKP$_{MP}$ (*pfkA-pfkB-pfpC*) in pHERD20T NcoI/HindIII | this study |
| pHERD20T-*pfkA*$^{GGM}$-*pfkB* | Ap$^R$, the 23$^{th}$-25$^{th}$ aa codon GGAGGCATG in *pfkA* was mutated to GCAGCGTGT in pHERD20T-*pfkA-pfkB* | this study |
| pHERD20T-*pfkA*$^{HRD}$-*pfkB* | Ap$^R$, the 131$^{th}$-133$^{th}$ aa codon CATCGCGAC in *pfkA* was mutated to TTCGAAAAT in pHERD20T-*pfkA-pfkB* | this study |
| pHERD20T-*pfkA*$^{DFG}$-*pfkB* | Ap$^R$, the 154$^{th}$-157$^{th}$ aa codon GACTTCGGT in *pfkA* was mutated to AATTATGCA in pHERD20T-*pfkA-pfkB* | this study |
| pHERD20T-*pfkA-pfkB*$^{GGM}$ | Ap$^R$, the 16$^{th}$-18$^{th}$ aa codon GGCGGCATG in *pfkB* was mutated to GCAGCGTGT in pHERD20T-*pfkA-pfkB* | this study |
| pHERD20T-*pfkA-pfkB*$^{HRD}$ | Ap$^R$, the 121$^{th}$-123$^{th}$ aa codon CATAGAGAT in *pfkB* was mutated to TTCGAAAAT in pHERD20T-*pfkA-pfkB* | this study |
| pHERD20T-*pfkA-pfkB*$^{DFG}$ | Ap$^R$, the 141$^{th}$-143$^{th}$ aa codon GACTTTGGC in *pfkB* was mutated to AATTATGCA in pHERD20T-*pfkA-pfkB* | this study |
| pHERD20T-*2208* | Ap$^R$, *2208* in pHERD20T NcoI/HindIII | this study |
| pHERD20T-*2209* | Ap$^R$, *2209* in pHERD20T NcoI/HindIII | this study |
| pHERD20T-*2210* | Ap$^R$, *2210* in pHERD20T NcoI/HindIII | this study |
| pHERD20T-*2208-2209* | Ap$^R$, *2208-2209* in pHERD20T NcoI/HindIII | this study |
| pHERD20T-*2208-2210* | Ap$^R$, *2208-2210* in pHERD20T NcoI/HindIII | this study |
| pHERD20T-*2209-2210* | Ap$^R$, *2209-2210* in pHERD20T NcoI/HindIII | this study |
| pHERD20T-*2208-2209-2210* | Ap$^R$, *2208-2209-2210* (KKP$_{SW}$ in marine *Shewanella* W3-18-1) in pHERD20T NcoI/HindIII | this study |
| pHERD20T-K$^{GFA}$-KP$_{SW}$ | Ap$^R$, *2208*$^{GFA}$*-2209-2210* (K$^{GFA}$KP$_{SW}$ with GFA mutation in marine *Shewanella* W3-18-1) in pHERD20T NcoI/HindIII | this study |
| pHERD20T-K$^{HRD+DFG}$-K$^{HRD+DFG}$P$_{SW}$ | ApR, 2208-2209$^{HRD+DFG}$-2210$^{HRD+DFG}$ (K$^{HRD+DFG}$K$^{HRD+DFG}$P$_{SW}$ with GFA mutation in marine *Shewanella* W3-18-1) in pHERD20T NcoI/HindIII | this study |
| pHERD20T-K$^{GFI}$-KP$_{EC039}$ | ApR, GFI mutation in *E. coli* 15EC039 KKP$_{EC039}$ (K$^{GFI}$KP$_{EC039}$) in pHERD20T NcoI/HindIII | this study |

**Table. S9** Oligonucleotides used for gene knockout and DNA sequencing. F indicates forward primer and R indicates reverse primer. The red letters indicate the mutation of nucleotides.

| Primer name | Sequence (5'-3') | Purpose |
|---|---|---|
| **Primers used for construction of MP-ΔPf6core and MP-ΔPf4ΔPf6core** | | |
| Pf6core-up-F | acgacggccagtgccaagcttTCTAAGTTCACATTCATGCACTCTTTC | |
| Pf6core-up-R | tgagcgggtctcaTTAGTCACCGTCGTAATCCCCC | Construction of pEX18Gm-ΔPf6core-up-down |
| Pf6core-down-F | tgactaaTGAGACCCGCTCAATGGACA | |
| Pf6core-down-R | ggtacccggggatcctctagaGAGTGGTGATAGCGAAACTCGAT | |
| Pf6core-conf-SF | TGAATCATCGCTACGCTCCTC | |
| Pf6core-conf-SR | ACGGCTGCTTGGTTGGTCT | Verification of Pf6core knockout in MP-ΔPf6core and MP-ΔPf4ΔPf6core |
| Pf6core-conf-LF | CATGGCAGACTCTATCAGCCA | |
| Pf6core-conf-LR | ATCGCATATCCCTTGCGCAGATA | |
| **Primers used for construction of pVIII::GFP and MP-pVIII::GFP** | | |
| *pVIII-gfp*-up-F | gccagtgccaagcttgcatgcAGGACAAACAAGACAAGACCCCG | |
| *pVIII-gfp*-up-R | TTCTCCTTTACTCGCCTTGCGCAACATGCT | |
| *pVIII-gfp*-F2 | GCAAGGCGAGTAAAGGAGAAGAACTTTTCACTGGA | Construction of pEX18Gm-pVIII::GFP-up-down |
| *pVIII-gfp*-R2 | CCAGAGCACCCGTTATTTGTATAGTTCATCCATGCCATG | |
| *pVIII-gfp*-down-F | ACAAATAACGGGTGCTCTGGTCGGTG | |
| *pVIII-gfp*-down-R | tatgaccatgattacgaattcCGGTGCCCTTGAGGATGTAA | |
| *pVIII-gfp*-conf-SF | GCGAATACAACATCGAGCCG | |
| *pVIII-gfp*-conf-SR | GGTCTGGGAGAAGGTGTAGGAAT | Verification of strains pVIII::GFP and MP-pVIII::GFP |
| *pVIII-gfp*-conf-LF | ATGAACATGTTTGCAACCCAA | |
| *pVIII-gfp*-conf-LR | TCAGTCCTTCAGCAGAATGAG | |
| **Primers used for construction of Δ*pilC* and PAO1::*pilC*$_{MP}$** | | |
| *pilC-up*-F | acgacggccagtgccaagcttTCTGCTCGTCTCAAGGTAAT | |
| *pilC-up*-R | ATGGCTGGCCAGGTAGTCGAGGAGGGGCATGGATTAATCCTTGGTCACGCGGTTGACTT | Construction of pEX18Gm-up-*pilC*-down, and pEX18Gm-up-*pilC*$_{MP}$-down |
| *pilC-down*-F | AAGTCAACCGCGTGACCAAGGATTAATCCATGCCCCTCCTCGACTACCTGGCCAGCCAT | |
| *pilC-down*-R | tatgaccatgattacgaattcAGTTGGTGATCGGCATCGAT | |
| *pilC*-conf-SF | GAGCAAGCCCGCAAAGAAG | Verification of strains Δ*pilC,* and PAO1::*pilC*$_{MP}$ |
| *pilC*-conf-SR | CCAGTTGCGCTCCATCATCT | |
| *pilC*-conf-LF | AAGATGCTGCTGGATGCCAT | |
| *pilC*-conf-LR | GATCAGCGGGTGCAGCAATT | |

**Primers used for construction of PAO1::*pilC*$_{MP}$::KKP$_{MP}$ and PAO1::*pilC*$_{MP}$::K$^{GFI}$KP$_{MP}$**

| | | |
|---|---|---|
| KKP$_{MP}$-F1 | acgacggccagtgccaagcttTAGTTGATTACCACTGAGGAGGTGG | Amplification of the 697 bp downstream of *pfkA* |
| KKP$_{MP}$-R1 | CGACCCAAGTACCGCCACCTAAATGGCACTCTAATAAATAAAATTAAATTTATCGCTTC | |
| KKP$_{MP}$-F2 | GAAGCGATAAATTTAATTTTATTTATTAGAGTGCCATTTAGGTGGCGGTACTTGGGTCG | Amplification of Gm resistance gene with its 303 bp promoter |
| KKP$_{MP}$-R2 | AGCACATCCTGTTATGATCCTTGATGAGTAGAGTTCCTATACTTTCTAGAGAATAGGAA | |
| KKP$_{MP}$-F3 | TTCCTATTCTCTAGAAAGTATAGGAACTCTACTCATCAAGGATCATAACAGGATGTGCT | Amplification of KKP cascades and upstream 404 bp promoter the start codon of *pf6r* was mutated to TGA) |
| KKP$_{MP}$-R3 | TGTCCCAGGTTCAAGTCCCGGTGTAGCCACCATGAGAATCAATACAGAGCAGAGCTAGG | |
| KKP$_{MP}$-F4 | CCTAGCTCTGCTCTGTATTGATTCTCATGGTGGCTACACCGGGACTTGAACCTGGGACA | Amplification of the 836 bp from Pf6 *attR* to upstream |
| KKP$_{MP}$-R4 | tatgaccatgattacgaattcGGGCAAGCCGGCGCGTTC | |
| KK$^{GFI}$P$_{MP}$-R3 | GAAAATTTCTTTCCCCATCAAGGATGACTAGGAATGAGCACAATACCAGACCGATATGA | Amplification of *pfkA*$^{GFI}$-*pfkB* from pHERD20T-*pfkA*$^{GFI}$-*pfkB* |
| KK$^{GFI}$P$_{MP}$-F4 | TCATATCGGTCTGGTATTGTGCTCATTCCTAGTCATCCTTGATGGGGAAAGAAATTTTC | Amplification of *pfpC* and the 836 bp from Pf6 *attR* to upstream with KKP$_{MP}$-R4, the template is PAO1::*pilC*$_{MP}$::KKP$_{MP}$ genomic DNA |

**Primers used for construction of pHERD20T-*mvaU* and related mutants**

| | | |
|---|---|---|
| pHERD20T-*mvaU*-F | aagaaggagatatacataccATGTCCAAACTTGCCGAGTTC | |
| pHERD20T-*mvaU*-R | cgacggccagtgccaagcttTTAGCGTTGCAGCCAGGATTC | |
| pHERD20T-*mvaU*$^{S2A}$-F | aagaaggagatatacataccATGGCCAAACTTGCCGAGTTC | |
| pHERD20T-*mvaU*$^{S2D}$-F | aagaaggagatatacataccATGGACAAACTTGCCGAGTTC | |
| pHERD20T-*mvaU*$^{S26A}$-F | GCTGAAAGCCGACAGCAGCCTGAAGCAGGAACTGGAAT | |
| pHERD20T-*mvaU*$^{S26A}$-R | GCTGCTGTCGGCTTTCAGCTTTTCCAGCAGGGCCAGTT | |
| pHERD20T-*mvaU*$^{S26D}$-F | AAGCTGAAAGACGACAGCAGCCTGAAGCAGGAACTGGAAT | |
| pHERD20T-*mvaU*$^{S26D}$-R | CAGGCTGCTGTCGTCTTTCAGCTTTTCCAGCAGGGCCAGT | |
| pHERD20T-*mvaU*$^{T50A}$-F | TACGGCATGGCCCTGCACAACATCATCGCCATCCTCGACC | |
| pHERD20T-*mvaU*$^{T50A}$-R | GTTGTGCAGGGCCATGCCGTACTTGTCCATCAACGCCTGC | |
| pHERD20T-*mvaU*$^{T50D}$-F | TACGGCATGGACCTGCACAACATCATCGCCATCCTCGACC | |
| pHERD20T-*mvaU*$^{T50D}$-R | GTTGTGCAGGTCCATGCCGTACTTGTCCATCAACGCCTGC | |
| pHERD20T-*mvaU*$^{T65A}$-F | AAGGCTCCGGTCGCCGTCAGCGCCGCTCCGCAGCGCCGTGCCCGC | |
| pHERD20T-*mvaU*$^{T65A}$-R | GGCGCTGACGGCGACCGGAGCCTTGGGGTCGAGGATGGCGATGAT | Construction of pHERD20T-*mvaU* and related single site mutations in MvaU |
| pHERD20T-*mvaU*$^{T65D}$-F | GCTCCGGTCGACGTCAGCGCCGCTCCGCAGCGCCGTGCC | |

| | | |
|---|---|---|
| pHERD20T-*mvaU*$^{T65D}$-R | GCTGACGTCGACCGGAGCCTTGGGGTCGAGGATGGCGAT | |
| pHERD20T-*mvaU*$^{S67A}$-F | AAGGCTCCGGTCACCGTCGCCGCCGCTCCGCAGCGCCGTGCCCGC | |
| pHERD20T-*mvaU*$^{S67A}$-R | GGCGGCGACGGTGACCGGAGCCTTGGGGTCGAGGATGGCGATGAT | |
| pHERD20T-*mvaU*$^{S67D}$-F | GCTCCGGTCACCGTCGACGCCGCTCCGCAGCGCCGTGCC | |
| pHERD20T-*mvaU*$^{S67D}$-R | GGAGCGGCGTCGACGGTGACCGGAGCCTTGGGGTCGAGGATGGCGAT | |
| pHERD20T-*mvaU*$^{S108A}$-R | cgacggccagtgccaagcttTTAGCGTTGCAGCCAGGATTCGACGGTTTCGGAACCGGCCTGTTCTTTC | |
| pHERD20T-*mvaU*$^{S108D}$-R | cgacggccagtgccaagcttTTAGCGTTGCAGCCAGGATTCGACGGTTTCGGAACCGTCCTGTTCTTTC | |
| pHERD20T-*mvaU*$^{S113A}$-R | cgacggccagtgccaagcttTTAGCGTTGCAGCCAGGCTTCGACGGTTTC | |
| pHERD20T-*mvaU*$^{S113D}$-R | cgacggccagtgccaagcttTTAGCGTTGCAGCCAGTCTTCGACGGTTTC | |
| **Primers used for cloning KKP$_{MP}$ components and domains of PfkA and PfkB** | | |
| pHERD20T-*pfkA*-F | aggagatatacatacccatgACGACTTCGAGAATCGGAAAGAC | Construction of pHERD20T-based constructions for expression of MPAO1 KKP components |
| pHERD20T-*pfkA*-R | acgacggccagtgccaagcttCTACTCATCAAGGATCATAACAGGATG | |
| pHERD20T-*pfkB*-F | aggagatatacatacccatgAGCACAATACCAGACCGATATGAATT | |
| pHERD20T-*pfkB*-R | acgacggccagtgccaagcttTCATAAGTTTATCTCCGGATGAGATAG | |
| pHERD20T-*pfpC*-F | aggagatatacatacccatgAATGTGAACTTAGAGAACGATATTACATTTC | |
| pHERD20T-*pfpC*-R | acgacggccagtgccaagcttCTAGTCATCCTTGATGGGGAAAGA | |
| *pfkA*$^{CSD}$-R | acgacggccagtgccaagcttTCAACATAAAAGTTCACATTTTTTTACC | Construction of pHERD20T-*pfkA*$^{CSD}$-*pfkB*, pHERD20T-*pfkA*-*pfkB*$^{FHA}$ and pHERD20T-*pfkA*$^{CSD}$-*pfkB*$^{FHA}$ together with primers pHERD20T-*pfkB*-F and pHERD20T-*pfkA*-R |
| *pfkB*$^{FHA}$-R1 | TTAGCGCTTGATGTCTGTTTTCTAACTATATGTGCTTCTCAAGTGTCTCTTTGATTGAC | |
| *pfkB*$^{FHA}$-F2 | GTCAATCAAAGAGACACTTGAGAAGCACATATAGTTAGAAAACAGACATCAAGCGCTAA | |
| pHERD20T-*pfkA*$^{CSD}$-R | acgacggccagtgccaagcttTCATATGTGCTTCTCAAGTGTCTCTTT | Construction of pHERD20T-*pfkA*$^{CSD}$, pHERD20T-*pfkA*$^{CSD}$, pHERD20T-*pfkB*$^{FHA}$ and pHERD20T-*FHA* together with primers pHERD20T-*pfkB*-F/R and pHERD20T-*pfkA*-F/R |
| pHERD20T-*CSD*-R | aggagatatacatacccATGTTAGAAAACAGACATCAAGCGC | |
| pHERD20T-*pfkB*$^{FHA}$-R | acgacggccagtgccaagcttCTAACATAAAAGTTCACATTTTTTTACCA | |
| pHERD20T-*FHA*-R | aggagatatacatacccATGTATGCAAAAAACCCAAGATATATTG | |
| **Primers used for construction of conserved domains in pHERD20T-*pfkA-pfkB*** | | |
| pHERD20T-*pfkA*$^{GGM}$-*pfkB*-F | ATCTCGATATTTAATCGAGGACTTAATCGGCGAAGCAGCGTGTCAATATGTTTACCG | F primers/HERD20T-*pfkA*-R and pHERD20T-*pfkB*-F/R primers were used to amplify the second and first fragments separately, the two fragments were fused using pHERD20T-*pfkB*-F and pHERD20T-*pfkA*-R and ligated into pHERD20T plasmid. |
| pHERD20T-*pfkA*$^{GGM}$-*pfkB*-R | CGGTAAACATATTGACACGCTGCTTCGCCGATTAAGTCCTCGATTAAATATCGAGAT | |
| pHERD20T-*pfkA*$^{HRD}$-*pfkB*-F | TGCCGGAGTAGTTTTCGAAAATCTGAAACCAAC | |
| pHERD20T-*pfkA*$^{HRD}$-*pfkB*-R | GTTGGTTTCAGATTTTCGAAAACTACTCCGGCA | |
| pHERD20T-*pfkA*$^{DFG}$-*pfkB*-F | GTTAAAAATTACTAATTATGCAATTGCAAAAATG | |
| pHERD20T-*pfkA*$^{DFG}$-*pfkB*-R | CATTTTTGCAATTGCATAATTAGTAATTTTTAAC | |

| | | |
|---|---|---|
| pHERD20T-*pfkA*-*pfkB*$^{GGM}$-F | TTGATGCAGCGTTAGGTACAATCTTCGTCTGC | |
| pHERD20T-*pfkA*-*pfkB*$^{GGM}$-R | GCAGACGAAGATTGTACCTAACGCTGCATCAA | |
| pHERD20T-*pfkA*-*pfkB*$^{HRD}$-F | GTAAACATAATTTTCGAAAATATCAAACCAAACAAC | |
| pHERD20T-*pfkA*-*pfkB*$^{HRD}$-R | GTTGTTTGGTTTGATATTTTCGAAAATTATGTTTAC | |
| pHERD20T-*pfkA*-*pfkB*$^{DFG}$-F | ATCATTAAAATATTTAATTATGCATTAGCAAGACACAC | |
| pHERD20T-*pfkA*-*pfkB*$^{DFG}$-R | GTGTGTCTTGCTAATGCATAATTAAATATTTTAATGAT | |
| *pfkA*-*pfkB*-linker-F | TTACGTTCGACCTATCTCATCCGGAGATAAACTTATGACGACTTCGAGAATCGGAAAGACTATA | Construction the double GGM, HDR, DFG mutations, the *pfkB* and *pfkA* single mutants were amplified using the above PCR products as templates, and fused with the primers and ligated into pHERD20T plasmid. |
| *pfkA*-*pfkB* -linker-R | TATAGTCTTTCCGATTCTCGAAGTCGTCATAAGTTTATCTCCGGATGAGATAGGTCGAACGTAA | |
| *pfkB*-in-R1 | TAAGTCATTGCCGTAGATATATTCTTGTACGATGCCAAGCCCGCGTTCG | Construction of pHERD20T-*pfkA*$^{GGM+HRD+DFG}$-*pfkB*, pHERD20T-*pfkA*-*pfkB*$^{GGM+HRD+DFG}$ and pHERD20T-*pfkA*$^{GGM+HRD+DFG}$-*pfkB*$^{GGM+HRD+DFG}$, mutant *pfkB* and *pfkA* with were amplified using above mutants as templates, and fused with these primers, and *pfkA*-*pfkB*-linker-F/R, then ligated into pHERD20T plasmid. |
| *pfkB*-in-F1 | CGAACGCGGGCTTGGCATCGTACAAGAATATATCTACGGCAATGACTTA | |
| *pfkB*-in-R2 | AATATTTTAATGATGCTTTCATGGTCTAGCATCATGTTGTTTGGTTTGA | |
| *pfkB*-in-F2 | TCAAACCAAACAACATGATGCTAGACCATGAAAGCATCATTAAAATATT | |
| *pfkA*-in-R1 | GGCTACGTTATGATGATTGACTTTGGCTGCTACAATGGCACTACGCCG | |
| *pfkA*-in-F1 | CGGCGTAGTGCCATTGTAGCAGCCAAAGTCAATCATCATAACGTAGCC | |
| *pfkA*-in-R2 | AACTCATTTAGTGAATATCCGCCAGAGATCATAACGTTTGTTGGTTTCA | |
| *pfkA*-in-F2 | TGAAACCAACAAACGTTATGATCTCTGGCGGATATTCACTAAATGAGTT | |
| **Primers used for cloning *Shewanella* W3-18-1 KKP$_{SW}$ components and mutants** | | |
| pHERD20T-*2208*-F | aggagatatacatacccatgTTTCAAAGGCTATTGCAAAAA | Construction of pHERD20T-based constructions for expression of KKP$_{SW}$ components and mutants |
| pHERD20T-*2208*-R | acgacggccagtgccaagcttCTACTTGGAATCATCATCTTCAACCT | |
| pHERD20T-*2209*-F | aggagatatacatacccatgATGATTCCAAGTAGATATGAACTCTG | |
| pHERD20T-*2209*-R | acgacggccagtgccaagcttTCATGAAACCACCTCTGGGTTAG | |
| pHERD20T-*2210*-F | aggagatatacatacccatgATCACATCGAATACTCTGATCG | |
| pHERD20T-*2210*-R | acgacggccagtgccaagcttTTAATTTAAGATAATGACTGGGTGAGC | |
| pHERD20T-*2210*$^{GFA}$-F | GCTATGTTGAGGGTTTGAAGCATAGTCACTATATCCGTTTTGAAT | |
| pHERD20T-*2210*$^{GFA}$-R | ATTCAAAACGGATATAGTGACTATGCTTCAAACCCTCAACATAGC | |
| pHERD20T-*2210*$^{HRD}$-F | CACACATTCAAGGAGTAATATTCGAAAATTTGAAGCCAAGTAATA | |
| pHERD20T-*2210*$^{HRD}$-R | TATTACTTGGCTTCAAATTTTCGAATATTACTCCTTGAATGTGTG | |
| pHERD20T-*2210*$^{DFG}$-F | TCGTCGGTAAAAGTTGCAATTGCATAATTGGTAATCTTTAGCTCT | |
| pHERD20T-*2210*$^{DFG}$-R | AGAGCTAAAGATTACCAATTATGCAATTGCAACTTTTACCGACGA | |

| | | |
|---|---|---|
| pHERD20T-*2209*$^{HRD}$-F | TACATAAGGCAGGAATTATTTTCGAAAATATCAAACCAAACAATA | |
| pHERD20T-*2209*$^{HRD}$-R | TATTGTTTGGTTTGATATTTTCGAAAATAATTCCTGCCTTATGTA | |
| pHERD20T-*2209*$^{DFG}$-F | CGGAGTTATCAAGGTCTTTAATTATGCACTTTCGAGAAAACTTGA | |
| pHERD20T-*2209*$^{DFG}$-R | TCAAGTTTTCTCGAAAGTGCATAATTAAAGACCTTGATAACTCCG | |
| **Primers used for cloning KKP components from other bacteria** | | |
| pHERD20T-*RS13785*-F | aggagatatacatacccatgCAAGATATATTAACAAAAAGGCTAA | |
| pHERD20T-*RS13795*-R | acgacggccagtgccaagcttTTATTTACAGCGAATAATTGGAAATGC | |
| pHERD20T-*STY4824*-F | aggagatatacatacccatgTTTACAGAACGACTTGCTCGC | |
| pHERD20T-*STY4822*-R | acgacggccagtgccaagcttTCAATCTTTCCAAAGAATTACAGGG | Construction of GFI mutants in PfkA homogues in the synthesized pHERD20T-KKP plasmids with different resources. |
| pHERD20T-*RS23220*-F | aggagatatacatacccatgAGCCGTGATTCTTACGAAAT | |
| pHERD20T-*RS23210*-R | acgacggccagtgccaagcttCTACTTAGCCTTGATAACTGGATGAGC | |
| pHERD20T-*3792*-F | aggagatatacatacccatgAGCCGTGATTCTTACGAAAT | |
| pHERD20T-*3794*-R | acgacggccagtgccaagcttCTACTTAGCCTTGATAACTGGATGAGC | |
| K$^{GFI}$KP$_{EC039}$-F | GATACTAGACTAGCTTATGACTATGCTCGTATACCAAACCAACCACAAG | |
| K$^{GFI}$KP$_{EC039}$-R | CTTGTGGTTGGTTTGGTATACGAGCATAGTCATAAGCTAGTCTAGTATC | |
| **Plasmid verification primers** | | |
| pEX18Ap-F/pEX18Gm-F | AATCTTCTCTCATCCGCCAAAACA | |
| pEX18Ap-R/pEX18Gm-R | CGCCCAATACGCAAACCGCCTCTC | Verification of constructions used for gene knockout and expression |
| pHDRD20T-F | ATCGCAACTCTCTACTGTTTCT | |
| pHDRD20T-R | TGCAAGGCGATTAAGTTGGGT | |
| **QPCR primers** | | |
| *pfiA*-qF | ATGCGGCTGACCTGGATT | |
| *pfiA*-qR | TGAGCGAACCTCCTGGAAA | |
| *pfkA*-qF | CGACCTGAAACCAACAAACGT | |
| *pfkA*-qR | TTAAACCATTGCTGAAAGGGAA | Quantification of Pf4 and Pf6 phages in effluents and expression levels of KKP$_{MP}$ components in MPAO1 biofilm at different days |
| *pfkB*-qF | GAAATGAGGTGCGTCGGTAT | |
| *pfkB*-qR | ACAAGAATATATCTACGGCAATGAC | |
| *pfpC*-qF | GCCATGACAACTGCTCCACT | |
| *pfpC*-qR | TTTTCTATGCTCGCTTGCTTT | |
| *gyrB*-qF | CAAGTACGAAGGCGGTCTGAAG | |

| | |
|---|---|
| *gyrB*-qR | GCAGAGCAGGTTCTCGTTGAA |
| *16SrRNA*-qF | TGGTTCAGCAAGTTGGATGTG |
| *16SrRNA*-qR | GTTTGCTCCCCACGCTTTC |
